# The anatomical compartment defines distinct immune remodeling within human visceral adipose tissue during aging

**DOI:** 10.64898/2026.07.06.736672

**Authors:** Rodrigo Castaneda, Hye-In Sim, Narae Park, Minsun Yu, Hye-Jin Lee, Hyun Yong Jin, Hyun Jung Kim, Kye Jin Park, Bo-Yeong Jin, Hyun Kyu Song, Heeju Ryu, Cheolju Lee, Kanghyun Ryu, Youngmin Ko, Hye-Sung Jo, Yoon Park, Rafael T. Han

**Author notes:** Correspondence to Youngmin Ko, Hye-Sung Jo, Yoon Park and Rafael T. Han. Rafael T. Han, Korea Institute of Science and Technology, Hwarang-ro 14-gil 5, Seongbuk-gu, Seoul, 02792, Republic of Korea., X: @RafaelTaeho, Yoon Park, Korea Institute of Science and Technology, Hwarang-ro 14-gil 5, Seongbuk-gu, Seoul, 02792, Republic of Korea.; Hye-Sung Jo, Department of Surgery, Seoul National University College of Medicine, 101 Daeharkro, Jongro-gu, Seoul, 03080, Republic of Korea., Youngmin Ko, Division of Kidney and Pancreas Transplantation, Department of Surgery, Asan Medical Center, University of Ulsan College of Medicine, 88, Olympic-ro 43-gil, Songpa-gu, Seoul, Republic of Korea. equal contribution (Rodrigo Castaneda, Hye-In Sim).

## Abstract

Age-associated inflammation varies across tissues, but whether distinct visceral adipose tissue (VAT) depots undergo inflammatory remodeling in a depot-specific manner remains unclear. Here we profiled human peri-organ VAT, comparing kidney-associated fat (KF, from kidney transplantation donors) and gallbladder-associated fat (GBF, from asymptomatic cholecystectomy patients with incidental polyps), using single-cell RNA sequencing, flow cytometry, intracellular protein profiling and *in situ* analysis. GBF showed broad tissue-residency and granzyme K-associated remodeling across conventional, regulatory and innate-like lymphocyte compartments, whereas KF showed stronger B cell remodeling, greater myeloid representation and more compartmentalized changes within resident effector-like CD8^+^ T cell states. We further identified age-associated CD20^+^ T cells with features consistent with local B-T cell interaction and an antigen-experienced, granzyme K-associated inflammatory memory phenotype. *In situ* analysis revealed age-associated myeloid accumulation and crown-like structure remodeling, accompanied by distinct myeloid inflammatory programs in KF and GBF. Finally, depot-specific immune signatures associated with clinical indices of adjacent kidney and liver function. These findings indicate that age-associated distinct immune programs within peri-organ VAT depots track with local tissue context and the state of the adjacent organ.

## Introduction

Aging is accompanied by a chronic, low-grade, inflammatory state that contributes to systemic dysfunction, including metabolic diseases. Comprehensive profiling in rodents has demonstrated that aging affects individual organ systems disproportionately, and has identified the immune compartment of adipose tissue as one of the most profoundly remodeled tissue niches during aging ^1,2^. Adipose tissue is a metabolically and immunologically active organ, secreting adipokines and both pro- and anti-inflammatory mediators into systemic circulation. It is broadly categorized into subcutaneous and visceral types. Visceral adipose tissue (VAT) accumulates with age and is more closely associated with metabolic dysfunction and disease than subcutaneous adipose tissue^3,4^. Studies in murine models have characterized remodeling of the VAT immune milieu during healthy aging, including the depletion of innate lymphoid cells such as natural killer cells and group 2 innate lymphoid cells (ILC2s), and the accumulation of several subtypes of T cells including GZMK^+^ T cells and fat-resident regulatory T cells^5–8^. In humans, however, comparable analyses have largely been confined to subcutaneous adipose tissue^9^ or to obesity and other metabolic diseases^10,11^, leaving normal VAT aging poorly defined.

VAT, moreover, is not anatomically or physiologically uniform. It comprises distinct depots that envelop different visceral organs, each embedded within a microenvironment defined by local vascular drainage and adjacent organ identity^3,12^. Intraperitoneal depots adjacent to digestive organs drain primarily through the portal venous system and lie in proximity to gut-associated barrier tissues, whereas retroperitoneal depots adjacent to organs such as the kidney lie within the systemic circulation. Whether such anatomical distinctions within the abdominal cavity translate into depot-specific patterns of age-associated immune remodeling in VAT, and whether these distinctive inflammatory programs reflect the metabolic function of the adjacent organ, remains incompletely understood in humans.

Here, we generated a multimodal immune atlas of human kidney-associated and gallbladder-associated visceral adipose tissue across age using single-cell RNA sequencing (scRNA-seq), flow cytometry, proteomics, and *in situ* analysis. We identified depot-specific age-associated changes across B cells, CD20-acquiring T cells, T_RM_ populations, Tregs and innate-like T cells, with several inflammatory and tissue residency-associated programs most evident in gallbladder-associated fat. These data provide a framework for understanding how anatomical location and adjacent organ context shape immune aging within human VAT.

## Results

### Differential immune landscape of aging human visceral adipose tissue (VAT) across anatomical sites

To determine the differential impact of aging on VAT immunity across distinct anatomical sites, we performed scRNA-seq on two different peri-organ fats. We collected peri-gallbladder fat (GBF) from asymptomatic patients with incidental findings of gallbladder polyps confirmed by pathology to be without acute cholecystitis or malignant lesions. Next, we obtained peri-renal fat (kidney fat; KF) from living donors undergoing nephrectomy for kidney transplantation. To minimize confounding from systemic disease and immunomodulatory exposures, donor inclusion was conducted under the following criteria: individuals were non-obese (18 < BMI < 30), had no history of prior abdominal surgery, systemic immunological or cardiovascular disorders, or active malignancy, and were not receiving medications known to alter immune function (e.g., systemic corticosteroids or immunosuppressants). Metabolic comorbidities such as hypertension (18/81), type 2 diabetes (1/81), and dyslipidemia (20/81) were uncommon in both cohorts (Detailed donor characteristics, including comorbidities, and medications are summarized in **Methods** and **Supplementary Table 1**). We then profiled the transcriptomes of FACS-sorted immune cells (CD45^+^) from GBF of 3 young (20-30 years old) and 4 old patients (65-87 years old), and KF of 4 young (20-30 years old) and 4 old patients (60-70 years old) (**Fig. 1a** and **Extended Data Fig. 1a,b**). The resulting quality-controlled and batch-corrected 96,341 cell events were visualized using uniform manifold approximation and projection (UMAP), which showed similar distributions across the two depots and other donor covariates including sex and BMI (**Fig. 1b** and **Extended Data Fig. 1c-e**). We identified 10 distinct clusters, each displaying canonical marker genes: resting T cells, CD4 effector T cells, CD8 effector T cells, proliferating T cells, NK cells, tissue-resident NK cells, dendritic cell (DC)-like myeloid cells, monocytes-macrophages, adipose-tissue macrophages (ATMs), and B cells (**Fig. 1b** and **Extended Data Fig. 1f,g**). Comparative analysis of cell frequency revealed a significant expansion of CD4 effector and CD8 effector T cells in both aged GBF and KF. In contrast, the proportion of NK lineage cells decreased with age in the KF, and myeloid lineages including ATMs and DC-like myeloid cells showed a relative decline in both depots (**Fig. 1c**). GBF samples were markedly enriched for B cells but KF samples were enriched for myeloid cell populations such as ATMs and DC-like myeloid cells (**Extended Data Fig. 1h**). Although we applied computational batch correction to integrate data from two independent medical centers^13^, technical heterogeneity in sample processing could confound cross-depot comparisons. To address this, we evaluated multiple sample-handling protocols and found that cryopreservation of dissociated single-cell suspensions best preserved cell viability and surface marker integrity (**Supplementary Fig. 1a**; see Methods for full protocol details).

**Figure 1.**
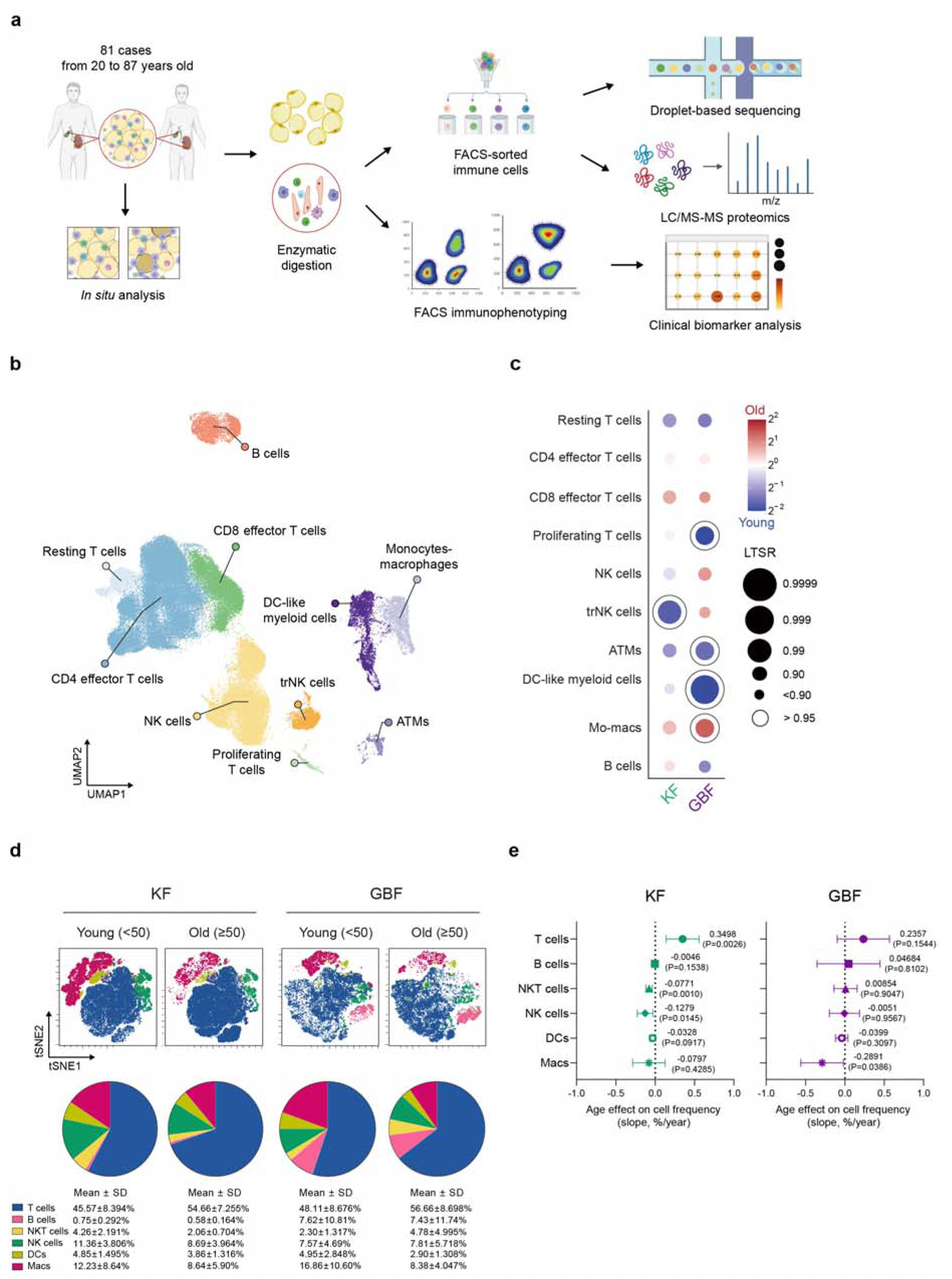
Differential immune landscape of aging human visceral adipose tissue (VAT) across anatomical sites. **a,** Schematic illustration of the study design with some graphical elements adopted from BioRender.com. **b**, Uniform manifold approximation and projection (UMAP) visualization of annotated cell events from the integrated VAT aging immune atlas. **c**, Bubble plots showing the differential abundance of cell clusters between age groups in the KF (first column) and GBF (second column). Local True Sign Rate (LTSR) denotes statistical confidence (ranging from 0 to 1), where 1 indicates a confident estimate. Color encodes log fold change between older and younger donors. Bubble size encodes LTSR; open circles indicate LTSR > 0.95. KF, kidney fat; GBF, gallbladder fat. **d**, t-SNE plots (upper) show the distribution of CD45^+^ immune cell subsets from young (<50 years) and old (≥50 years) donors in KF and GBF, as assessed by flow cytometry. Pie charts (lower) show the mean proportional composition of these subsets as percentages of total CD45^+^ cells in the corresponding group shown directly above, representing donor-level group means. In both panels, subsets are color-coded according to the key (lower left). t-SNE, t-distributed stochastic neighbor embedding; NKT, natural killer T cell; NK, natural killer cell; DC, dendritic cell. **e**, Forest plots showing the age-associated slope on the frequency of each immune cell subset, as assessed by flow cytometry. Each dot represents the estimated slope, expressed as percentage points per year, with 95% confidence intervals (CIs). Unless otherwise indicated, all analyses in Fig. 1 were performed using the same donor cohort (KF, n = 22; GBF, n = 22). For age-stratified analyses, donors were divided into young (<50 years) and old (≥50 years) groups, with n = 9-13 donors per age/depot group.

Using this standardized sample preparation, we analyzed an independent validation cohort to characterize age-related immunophenotypic shifts across the lifespan in the two depots. Consistent with our scRNA-seq data, flow cytometric analysis of VATs demonstrated that the frequency of T cells was positively correlated with increasing patient age in both GBF and KF (**Fig. 1d,e**, **Extended Data Fig. 1i**, and **Supplementary Fig. 1b**) while the frequency of innate-like lymphoid cells including NK cells were negatively correlated with age in KF (**Fig. 1d,e** and **Supplementary Fig. 1b&c**), further supporting our scRNA-seq analysis. We also observed a significantly higher proportion of B cells in the GBF than in the KF (**Supplementary Fig. 1b**). In contrast, the frequency of myeloid lineage cells was negatively correlated with the age of patients, more strongly in GBF in both sequencing (**Fig. 1c**) and flow cytometric analyses (**Fig. 1d,e** and **Supplementary Fig. 1b**).

This apparent age-dependent decline in myeloid frequency, however, likely reflects a compositional dilution by expanding lymphoid populations rather than an absolute reduction in myeloid cells. In addition, enzymatic dissociation, which can selectively deplete tissue-adherent myeloid populations, may further contribute to their under-representation^11,14^. We therefore performed complementary *in situ* analyses. Our immunohistochemical quantification of IBA1^+^ myeloid cells revealed an age-dependent increase in KF (r=0.63, *p*=0.0023) whereas the accumulation in the GBF showed a similar trend but did not reach statistical significance (r=0.32, *p*=0.1811) (**Extended Data Fig. 1j,k**). Taken together, our scRNA-seq dataset, along with flow cytometry and *in situ* analyses reveals depot-specific immune remodeling in GBF and KF during aging.

### Age-associated B cell remodeling and accumulation of CD20^+^ T cells in human VAT

To profile B cells in VATs, we first mapped their heterogeneity by scRNA-seq and identified transcriptionally distinct naïve-like/transitional, early, memory, activated, APC-like, plasma cell, age-associated plasma cell and age-associated B cell (ABC) states^15^ (**Fig. 2a** and **Extended Data Fig. 2a**). Using a B cell surface-marker FACS panel, we then classified CD19^+^ B cells into naïve-like/transitional, memory, plasmablast, double-negative (DN) and ABC compartments. ABCs were identified among CD19^+^CD20^+^CD27^-^ cells and distinguished from naïve-like/transitional cells by their surface-marker profile (**Fig. 2b**). Although minor numerical differences were observed, the overall distribution of major B cell subsets was broadly comparable between KF and GBF (**Fig. 2c** and **Extended Data Fig. 2b**). Modeling age as a continuous variable revealed tissue-dependent B cell remodeling (**Fig. 2d** and **Supplementary Fig. 2a**). In KF, the naïve-like/transitional compartment declined with age, whereas DN B cells and plasmablasts showed positive age-associated trends; ABCs showed a more modest association. GBF showed similar directional trends, but the redistribution of non-naïve compartments was less pronounced (**Fig. 2d** and **Extended Data Fig. 2c**). To further characterize age-associated transcriptional changes, we examined differentially expressed genes across B cell states. The age-associated plasma cell state was enriched for *NR4A1*, *NR4A2*, *NR4A3*, *JCHAIN*, *RAB20*, *CXCR5*, *CXCL8* and *IL1B* (**Fig. 2e** and DEG list in **Supplementary Table 2**). These findings highlight transcriptional features accompanying age-related B cell remodeling. We also stratified B cells by HLA-DR and IgGκ expression to examine surface-marker–defined activation and humoral states (**Fig. 2f,g**). Among total B cells, HLA-DR^+^ frequency decreased with age in KF, whereas IgGκ^+^ frequency showed no clear age association; corresponding trends in GBF were weaker (**Fig. 2f**). Subset-level HLA-DR/IgGκ analysis showed tissue-and subset-specific patterns rather than a uniform shift across the B cell compartment (**Fig. 2g** and **Supplementary Fig. 2b**). In KF, age-associated changes were more evident within plasmablast-, ABC-and DN-associated subsets, whereas GBF showed more limited changes across most HLA-DR/IgGκ-defined states. Together, these data show that age-associated B cell remodeling in VATs involves both compositional and surface-marker–defined state changes, with stronger remodeling in KF than in GBF. We also identified CD20^+^ T cells in VATs, including both CD4^+^ and CD8^+^ populations in KF and GBF (**Fig. 2h**). Because CD20 is not typically expressed by T cells^16^, we examined the scRNA-seq data for CD20 transcripts in T cells. CD20 transcripts were not detected in T cells (**Extended Data Fig. 1f**), suggesting that surface CD20 was not explained by endogenous transcription. We therefore examined whether CD20^+^ T cells carried B cell-associated surface molecules. CD86 was selectively detected on CD20^+^ T cells relative to CD20^-^ T cells (**Fig. 2i**), consistent with B–T cell contact–associated membrane acquisition. The frequency of CD20^+^ T cells showed positive age-associated trends in both KF and GBF (**Fig. 2j**).

**Figure 2.**
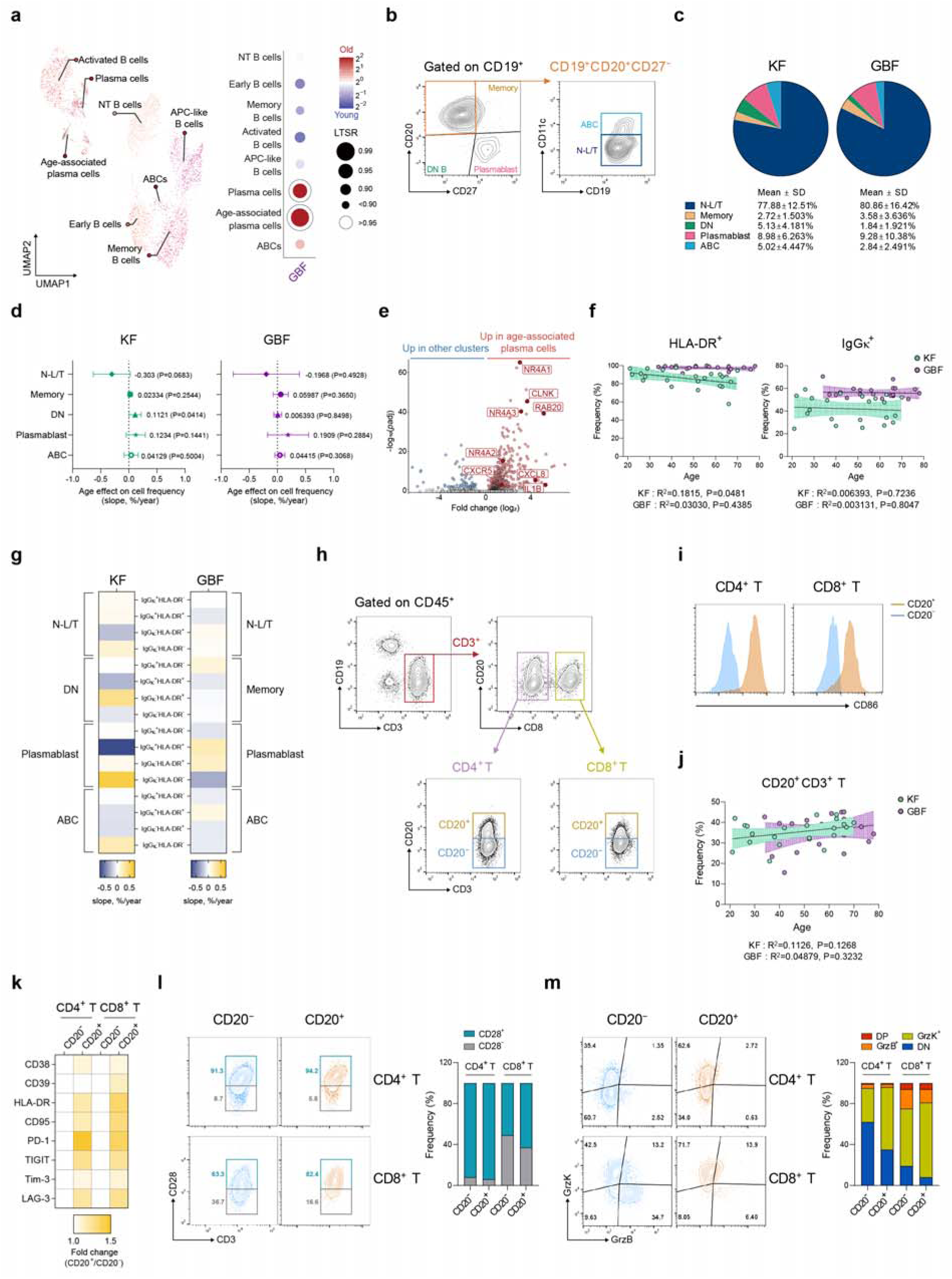
B cell remodeling and CD20^+^ T cell enrichment with age in VAT. **a**, UMAP of B cells from scRNA-seq, colored by subcluster (left) : naïve/transitional (NT), early, memory, activated, and antigen-presenting-cell-like (APC-like) B cells, plasma cells, age-associated plasma cells, and age-associated B cells (ABCs). Bubble plot of the relative abundance of each B cell subcluster in older versus younger GBF donors (right). Color encodes log_2_ fold change (old/young); bubble size encodes LTSR; open circles indicate LTSR > 0.95. **b,** Representative flow cytometry gating plots used to define CD19^+^ B cell subsets. **c**, Pie charts showing the mean proportional composition of the B cell subsets defined in b, expressed as percentages of total CD19^+^ cells in each age/depot group. **d**, Forest plots showing age-associated slopes in the frequencies of each B cell subset. Points indicate estimated slopes, expressed as percentage points per year; horizontal lines indicate 95% CIs. **e**, Volcano plot of differentially expressed genes in the age-associated plasma cell subcluster versus all other B cell subclusters. **f**, Regression graphs showing the frequency of HLA-DR^+^ (left) and IgGκ^+^(right) cells among CD19^+^ B cells against age. Lines indicate simple linear regression fits with 95% CIs (shaded). **g**, Heatmaps showing age-associated slopes in the frequencies of HLA-DR and IgGκ co-expression states within each B cell subset, estimated by simple linear regression and expressed as percentage points per year. **h**, Representative flow cytometry gating plots used to identify CD20^+^ T cells in KF. **i**, Representative histograms showing CD86 expression in CD20^+^ and CD20^-^CD4^+^ T cells and CD8^+^ T cells. **j**, Regression graphs showing the frequency of CD20^+^ T cells (CD20^+^CD3^+^) against age. Lines indicate simple linear regression fits with 95% CIs (shaded). **k**, Heatmap showing the fold change (CD20^+^/CD20^-^) in CD38, CD39, HLA-DR, CD95, PD-1, TIGIT, TIM-3 and LAG-3 expression in CD4^+^ and CD8^+^ T cells in KF (n = 7). **l**,**m**, Representative flow cytometry plots (upper) and bar graphs (lower) showing the frequency of CD28^+^ and CD28^-^ populations (**l**) and of GrzK/GrzB co-expression states (**m**) within CD20^-^ and CD20^+^ subsets of CD4^+^ and CD8^+^ T cells in KF (n = 7). Pie charts in **c** show the group mean composition; data in **l** and **m** are presented as mean ± s.d. *p* value in **d** were derived from simple linear regression. *R*² and *p* value in **f** and **j** were derived from simple linear regression.

To further characterize CD20^+^ T cells, we examined tissue-residency, activation and effector-associated markers. CD20^+^ and CD20^-^ T cells showed no major differences in CD69/CD103-defined tissue-resident memory composition (**Extended Data Fig. 2d**). However, CD20^+^ T cells were preferentially enriched for antigen experience/exhaustion-associated markers, including PD-1, compared with CD20^-^ T cells across both CD4^+^ and CD8^+^ compartments (**Fig. 2k and Extended Data Fig. 2e,f**). We next examined the CD28–granzyme axis, as CD28 loss is a hallmark of aged or senescent-like T cells, whereas *GZMK*^+^ T cells have been associated with inflammaging and inflamed tissue states^6,17–19^. In VATs, GrzK expression was preferentially associated with CD28^+^ cells, while GrzB was enriched in the CD28^-^ compartment (**Extended Data Fig. 2g**). CD20^+^ T cells contained a higher fraction of CD28^+^ cells and were skewed toward a GrzK^+^GrzB^-^ phenotype, whereas CD20^-^ T cells, particularly within the CD8^+^ compartment, were more enriched for GrzB-only cells (**Fig. 2l,m**). Thus, CD20^+^ T cells represent a surface CD20-acquiring, antigen-experienced T cell state with CD28-retaining and GrzK-associated features, rather than a CD28^-^GrzB^+^ terminal cytotoxic phenotype.

### T cell lineage composition and T_RM_-state heterogeneity across VAT depots and age

To define the lymphocyte landscape in VATs, we first used scRNA-seq to identify major T, NK and ILC populations, including conventional CD4^+^ and CD8^+^ T cell states, Tregs, MAIT cells, γδ T cells, NKT cells, TRNK cells, NK cells and ILCs (**Fig. 3a** and **Extended Data Fig. 3a**). FACS analysis using canonical surface markers further identified the corresponding major populations, including NKT cells, Vδ1^+^ and Vδ2^+^ γδ T cells, MAIT cells, CD4^+^ and CD8^+^ T cells, and CD4^+^ or CD8^+^ Tregs (**Fig. 3b**). Across KF and GBF, CD4^+^ and CD8^+^ T cells represented the predominant T cell lineages, whereas NKT, MAIT, γδ T and regulatory T cell populations comprised smaller fractions (**Fig. 3c**). The overall lineage distribution was broadly comparable between tissues, with minor differences in selected subsets (**Extended Data Fig. 3b**). When age was modeled continuously, most populations showed modest age-associated trends, with a decrease in NKT cells being more apparent in KF than in GBF (**Fig. 3d** and **Supplementary Fig. 3a**).

**Figure 3.**
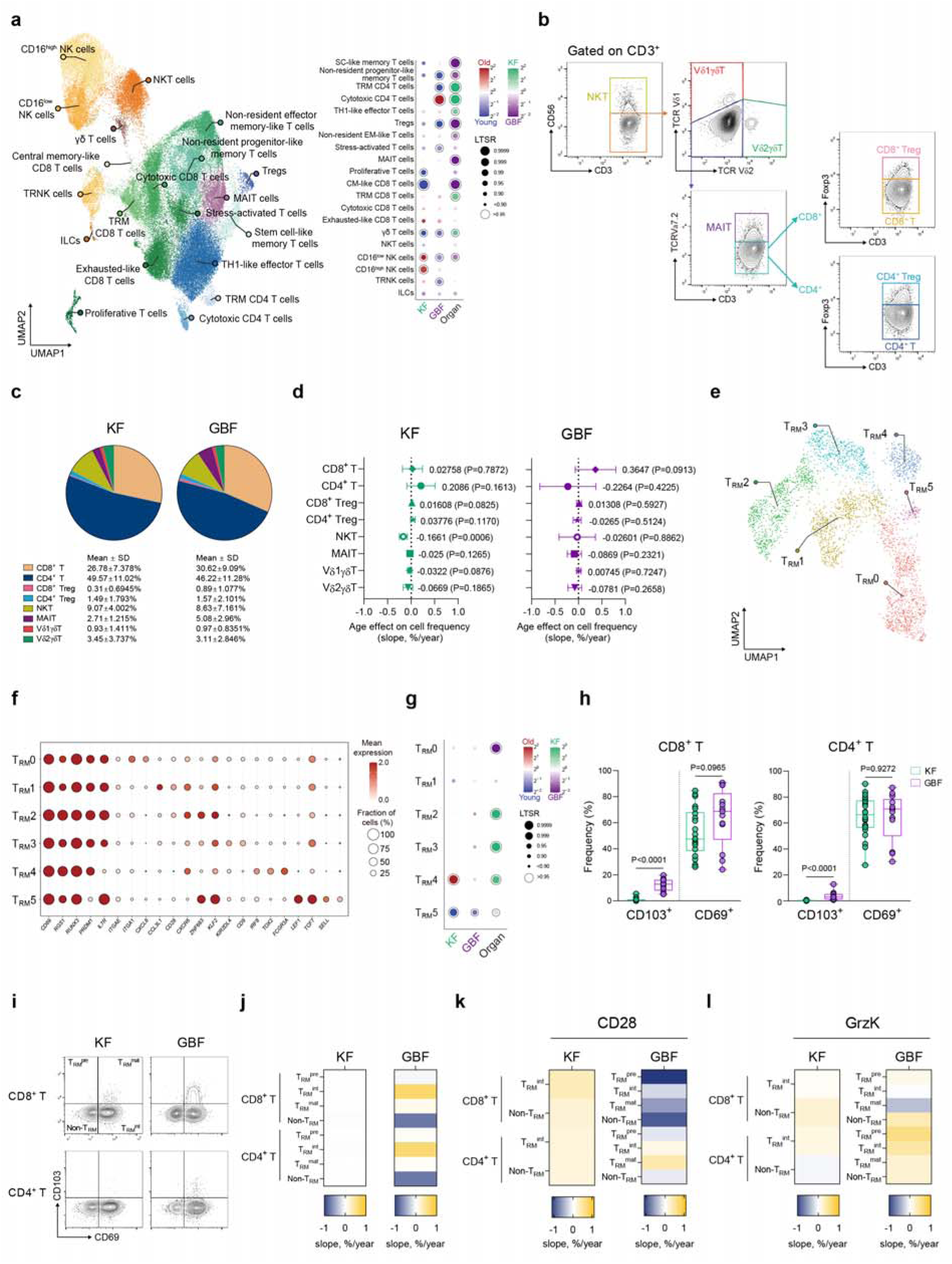
T cell subset distribution and T_RM_-state heterogeneity with age in VAT. **a**, Left: UMAP visualization of annotated non-B lymphoid cell clusters. Right: bubble plot of age- and depot-level differential abundance for each cluster. For the KF and GBF columns, color encodes log fold change between older and younger donors (red, old; blue, young); for the Organ column, color encodes log fold change between depots (green, KF; purple, GBF). Bubble size encodes the local true sign rate (LTSR), a measure of statistical confidence ranging from 0 to 1; open circles indicate LTSR > 0.95. **b**, Representative gating plots for flow cytometry analysis of CD3^+^ T cells. **c**, Pie charts showing the proportional composition of each T cell subset. **d**, Forest plots showing the age-associated effect on the frequency of each T cell subset. Each dot represents the slope (% per year) with its 95% CIs. **e,** UMAP visualization of the six tissue-resident memory T cell (T_RM_0–T_RM_5) subclusters. **f,** Dot plot of marker gene expression across the six T_RM_ subclusters. Dot size encodes the fraction of cells expressing each gene (%); color encodes mean scaled expression. **g,** Bubble plot of age- and depot-level differential abundance for each T_RM_ subcluster. Color encoding as in a; bubble size encodes LTSR, with open circles indicating LTSR > 0.95. **h**, Box plots showing the frequency of CD103^+^ and CD69^+^ cells within CD8^+^ T cells (left) and CD4^+^ T cells (right). **i**, Representative gating plots for flow cytometry analysis of CD69/CD103-defined T_RM_ subsets (T_RM_^pre^, T_RM_^int^, T_RM_^mat^ and Non-T_RM_) within CD8^+^ and CD4^+^ T cells. T_RM_^pre^, pre-T_RM_ cells; T_RM_^int^, intermediate-T_RM_ cells; T_RM_^mat^, mature-T_RM_ cells. **j–l**, Heatmaps showing the age-associated slope (% per year), derived from simple linear regression, of T_RM_ subset frequencies (**j**), CD28^+^ frequency within each T_RM_ subset (**k**) and GrzK^+^ frequency within each T_RM_ subset (**l**) in CD8^+^ and CD4^+^ T cells. Unless otherwise indicated, analyses in this figure used KF (n = 22) and GBF (n = 13) donors; the smaller GBF cohort reflects insufficient tissue for the T cell panel in the remaining GBF samples. Pie charts in **c** show the group mean composition. *p* value in **d** were derived from simple linear regression. *p* value in **h** were derived from two-tailed unpaired *t*-tests; box-plot centre lines indicate the median, box limits the interquartile range and whiskers the minimum to maximum.

We next focused on CD8^+^ T_RM_ cells, which formed a prominent resident T cell compartment in the scRNA-seq dataset. Subclustering identified six CD8^+^ T_RM_ states, designated T_RM_0–T_RM_5 (**Fig. 3e**). These states shared core residency-associated genes, including *CD69*, *RGS1*, *RUNX3* and *PRDM1*, but separated along retention-, memory- and effector-associated axes marked by *ITGAE*, *ITGA1*, *CXCR6*, *ZNF683*, *CD28*, *GZMK*, *LEF1*, *TCF7*, *SELL and S1PR1* (**Fig. 3f**). Among the tissue-biased states, GBF-enriched T_RM_0 showed a core resident profile without the stronger GZMK-, ZNF683- or effector-associated features observed in other clusters. In contrast, KF-enriched T_RM_2 and T_RM_3 expressed higher levels of *CXCR6*, *ZNF683* and *GZMK*, indicating a tissue-retention and GrzK-associated resident state. KF-enriched T_RM_4 showed a more differentiated effector-associated profile, with increased expression of genes such as *KLRD1*, *GNLY*, *IFNG*, *EOMES* and *CD244*. Age-associated changes were also cluster-selective: T_RM_4 was increased in old KF, whereas T_RM_5, marked by *IL7R* together with memory- or recirculation-associated genes including *TCF7*, *LEF1*, *SELL* and *S1PR1*, was reduced with age in both tissues (**Fig. 3e-g**, **Extended Data Fig. 3c**, and **Supplementary Table 3** for the full DEG list).

Building on the CD8^+^ T_RM_ transcriptional states, we next used FACS analysis to define residency-associated surface states and to examine aging-relevant phenotypic markers within those states. CD103 expression was markedly higher in GBF than in KF in both CD8^+^ and CD4^+^ T cells, whereas CD69 expression was broadly comparable between tissues (**Fig. 3h**). CD69 and CD103 were then used to define four surface-marker states^20^: CD69^-^CD103^-^Non-T_RM_, CD69^+^CD103^-^ T_RM_^int^, CD69^-^CD103^+^ T_RM_^pre^ and CD69^+^CD103^+^ T_RM_^mat^ cells (**Fig. 3i** and **Extended Data Fig. 3d**). In GBF, Non-T_RM_ cells decreased and T_RM_^int^ cells increased with age in both CD8^+^ and CD4^+^ T cells, whereas corresponding changes in KF were more limited (**Fig. 3j** and **Supplementary Fig. 3b**). Within these CD69/CD103-defined states, we next applied the CD28-GrzK axis defined above to assess whether aging altered memory/differentiation and inflammaging-associated phenotypes within T_RM_-related compartments. CD28^+^ frequencies were largely preserved across KF T_RM_ states but declined with age in selected GBF CD8^+^ compartments, most notably Non-T_RM_ and T_RM_^pre^ cells (**Fig. 3k, Extended Data Fig. 3e** and **Supplementary Fig. 3c**). In contrast, GrzK expression showed limited or inconsistent age-associated changes across KF T_RM_ states, whereas GBF showed an overall age-associated increase in GrzK^+^ frequencies across T cell compartments (**Fig. 3l, Extended Data Fig. 3f and Supplementary Fig. 3d**). Proteomic analysis of non-naïve CD4^+^ and CD8^+^ T cells from GBF, performed to capture intracellular changes not readily resolved by scRNA-seq or FACS, further showed age-related alterations in cytotoxic, mitochondrial and stress-response pathways (**Extended Data Fig. 3g–i** and **Supplementary Table 4** for individual protein abundance data). Together, these data suggest that aging promotes distinct forms of T_RM_-associated inflammaging across VAT depots. In GBF, aging was accompanied by broader T cell remodeling toward a differentiated CD28-low, GrzK-enriched phenotype, whereas the age-associated changes in KF were confined within resident effector-like CD8^+^ T_RM_ states, indicating a more compartmentalized T_RM_ response rather than broad inflammaging across the T cell pool.

### GBF-biased inflammatory adaptation of Tregs and innate-like T cells in aging VAT

To determine whether Tregs also showed depot-specific remodeling in VATs, we first separated CD4^+^Foxp3^+^ Tregs into naïve-like and memory subsets by FACS. CD45RO^+^ memory Tregs predominated in both KF and GBF, with GBF showing a modestly higher naïve-like fraction than KF (**Fig. 4a**). When age was modeled continuously, naïve-like and memory Treg frequencies showed reciprocal trends between depots: naïve-like Tregs tended to increase with age in KF but decrease in GBF, whereas memory Tregs showed the opposite pattern (**Fig. 4b**). Analysis of residency-associated surface markers in CD4^+^ Tregs showed markedly higher CD103 expression in GBF than in KF, with a more modest increase in CD69 (**Fig. 4c**). We therefore stratified memory Tregs into CD69^+^ and CD69^-^fractions to distinguish a CD69-associated tissue-resident regulatory population from a less tissue-retained comparator within the memory Treg compartment^21–23^. CD69^+^ memory Tregs were more frequent in GBF than in KF, consistent with the higher CD69 expression observed at the total Treg level (**Fig. 4d**). We then asked whether CD69-defined memory Tregs differed in aging- and inflammation-associated phenotypes. CD28 was retained at high frequency in most memory Treg subsets, but GBF CD69^-^ memory Tregs showed lower CD28 expression than their CD69^+^ counterparts. GrzK was included because GrzK is associated with inflammaging in conventional T cells and has also been reported in Tregs undergoing Th1-response-associated specialization^24^. CD69^+^ memory Tregs showed higher GrzK expression than CD69^-^ memory Tregs, particularly in GBF, whereas Ki-67 was more enriched in the CD69^-^ fraction (**Fig. 4e and Extended Data Fig. 4a**). Age-associated analysis further showed stronger GrzK induction in GBF memory Tregs, indicating that the GBF Treg compartment acquires an inflammation-adapted regulatory phenotype with age (**Fig. 4f** and **Supplementary Fig. 4a,b**).

**Figure 4.**
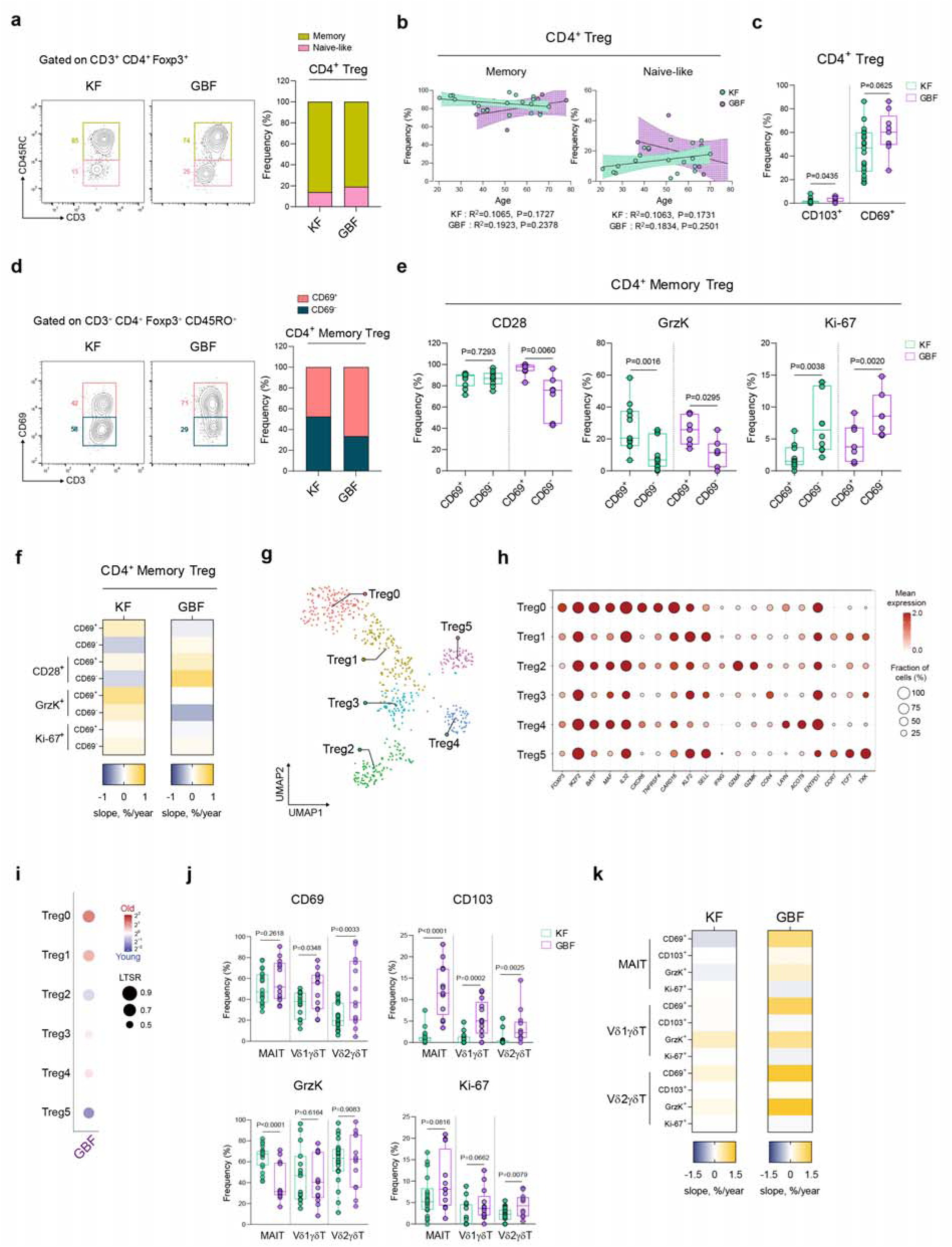
Age-associated phenotype of CD4^+^ Tregs and innate-like T cells in VAT. **a,** Representative flow cytometry plots (left) and bar graphs (right) showing the proportion of naïve-like (CD45RO^-^) and memory (CD45RO^+^) subsets within CD4^+^ Tregs. **b**, Regression graphs showing the frequency of memory (left) and naïve-like (right) CD4^+^ Tregs against age. Lines indicate simple linear regression fits with 95% CIs (shaded). **c**, Box plots showing the frequency of CD103^+^ and CD69^+^ cells within CD4^+^ Tregs. **d**, Representative plots (left) and bar graphs (right) showing the proportion of CD69^+^ and CD69^-^ subsets within CD4^+^ memory Tregs. **e**, Box plots showing the frequency of CD28^+^, GrzK^+^ and Ki-67^+^ cells within CD69^+^ and CD69^-^subsets of CD4^+^ memory Tregs. **f**, Heatmaps showing the age-associated slope (% per year), derived from simple linear regression, of CD69^+^ and CD69^-^ subset frequencies within CD4^+^ memory Tregs and of CD28^+^, GrzK^+^ and Ki-67^+^ frequencies within the CD69^+^ and CD69^-^subsets. **g**, UMAP visualization of Treg subclusters. **h**, Dot plot of marker gene expression across the six Treg subclusters. Dot size encodes the fraction of cells expressing each gene. Color encodes scaled mean expression. **i**, Bubble plot of age-associated differential abundance for each Treg subcluster in the GBF depot. Color encodes the log fold change in subcluster proportion between older and younger donors (red, enriched in aged; blue, enriched in young); bubble size encodes the local true sign rate (LTSR). **j**, Box plots showing the frequency of CD69^+^, CD103^+^, GrzK^+^ and Ki-67^+^ cells within MAIT cells, Vδ1^+^ γδ T cells and Vδ2^+^ γδ T cells. **k**, Heatmaps showing the age-associated slope (% per year), derived from simple linear regression, of CD69^+^, CD103^+^, GrzK^+^ and Ki-67^+^ frequencies within MAIT cells, Vδ1^+^ γδ T cells and Vδ2^+^ γδ T cells. Treg analyses (a–f) used KF (n = 19) and GBF (n = 9) donors, whereas innate-like T cell analyses (j,k) used KF (n = 22) and GBF (n = 13) donors. In both cell types the GBF cohort is smaller than the KF cohort because limited tissue availability precluded the full flow cytometry panel in some GBF donors. Data in **a** and **d** are presented as mean ± s.d. In box plots (**c**, **e**, **j**), centre lines indicate the median, box limits the interquartile range and whiskers the minimum to maximum. *R*² and *p* value in **b** were derived from simple linear regression. *p* value in **c** and **j** were derived from two-tailed unpaired *t*-tests; *p* value in **e** were derived from two-tailed paired *t*-tests.

To define Treg heterogeneity at the transcriptomic level, we subclustered Tregs from the scRNA-seq dataset and identified six states, designated Treg0–Treg5 (**Fig. 4g**). These clusters shared canonical Treg-associated genes, including *FOXP3* and *IKZF2*, but differed along activation, tissue-adaptation, cytotoxic/inflammatory and recirculation-associated axes marked by genes such as *BATF*, *MAF*, *IL32*, *CXCR6*, *GZMA*, *GZMK*, *ENTPD1*, *CCR7*, *TCF7* and *TXK* (**Fig. 4h**). Among these states, Treg0 showed an activated tissue-adapted profile, with expression of *FOXP3* and *IKZF2* together with *BATF*, *MAF*, *IL32*, *CXCR6* and *ENTPD1*. In GBF, aging was associated with enrichment of Treg0 and relative reduction of Treg5, a state marked by recirculation or memory-associated genes including *CCR7*, *TCF7* and *TXK* (**Fig. 4i**, **Extended Data Fig. 4b** and **Supplementary Table 5** for the full DEG list). Thus, consistent with the phenotyping data, GBF aging was associated with a shift toward tissue-retained, activated memory Treg phenotypes rather than a simple expansion of total Tregs.

We then examined whether residency-associated remodeling extended to innate-like T cell populations. MAIT cells and Vδ1^+^ or Vδ2^+^ γδ T cells showed higher CD103 expression in GBF than in KF, and CD69 was also higher in GBF, particularly within γδ T cell subsets (**Fig. 4j**). GrzK and Ki-67 showed more subset-specific differences, indicating that residency marker expression and effector/proliferative features were not uniformly coupled across innate-like populations. Age-associated analysis showed stronger positive slopes for CD69 and GrzK in GBF than in KF across MAIT and γδ T cell subsets, whereas KF showed weaker or mixed age-related changes (**Fig. 4k** and **Supplementary Fig. 4c**). Together, these data indicate that GBF is characterized by a broader tissue-residency and inflammaging-associated remodeling program across Tregs and innate-like T cells, while KF shows more limited or subset-restricted age-associated changes.

### Distinct inflammatory program of VAT myeloid cells during aging

To better understand cellular and molecular landscape of myeloid cells during aging, we clustered myeloid cells into ten heterogeneous subclasses of inflammatory myeloid cells, AT macrophages (ATMs), lipid-associated macrophages (LAMs), CD1C^high^ macrophages, *CCR2*^high^ myeloid cells, non-classical monocytes, neutrophils, conventional type 1 dendritic cells (cDC1s), conventional type 2 dendritic cells (cDC2s), and plasmacytoid dendritic cells (pDCs) based on differential expression of marker genes (**Fig. 5a** and **Extended Data Fig. 5a**). This taxonomy recovered macrophage states previously described in human white adipose tissue including *TREM2* and *CD9* expressing LAMs, *SELENOP* and *FOLR2* expressing ATMs, and inflammatory myeloid cell ^11,25^. *CCR2*^high^ myeloid cells expressing *VCAN*, indicative of recent trafficking from the blood, were also detected. However, this cluster was derived disproportionately from a single donor in our cohort (**Extended Data Fig. 5b**), suggesting inter-individual variation rather than depot- or age-related biology. We next compared myeloid composition between young and aged donors across the two depots. Overall myeloid composition was largely stable with age in both depots (**Fig. 5b**). LAMs were enriched in aged GBF, although their sparsity in KF precluded reliable quantification in that depot. Homeostatic ATMs were comparatively enriched in GBF (**Extended Data Fig. 5c**) and trended toward an age-associated increase in this depot, although this did not reach statistical significance (**Fig. 5b**). Consistent with the scRNA-seq data, flow cytometry with CD11c demarcating inflammatory macrophages^26^ demonstrated that the proportion of CD11c^-^CD206^+^ ATMs showed a trend toward an increase with age in GBF but not in KF (**Extended Data Fig. 5d-g**).

**Figure 5.**
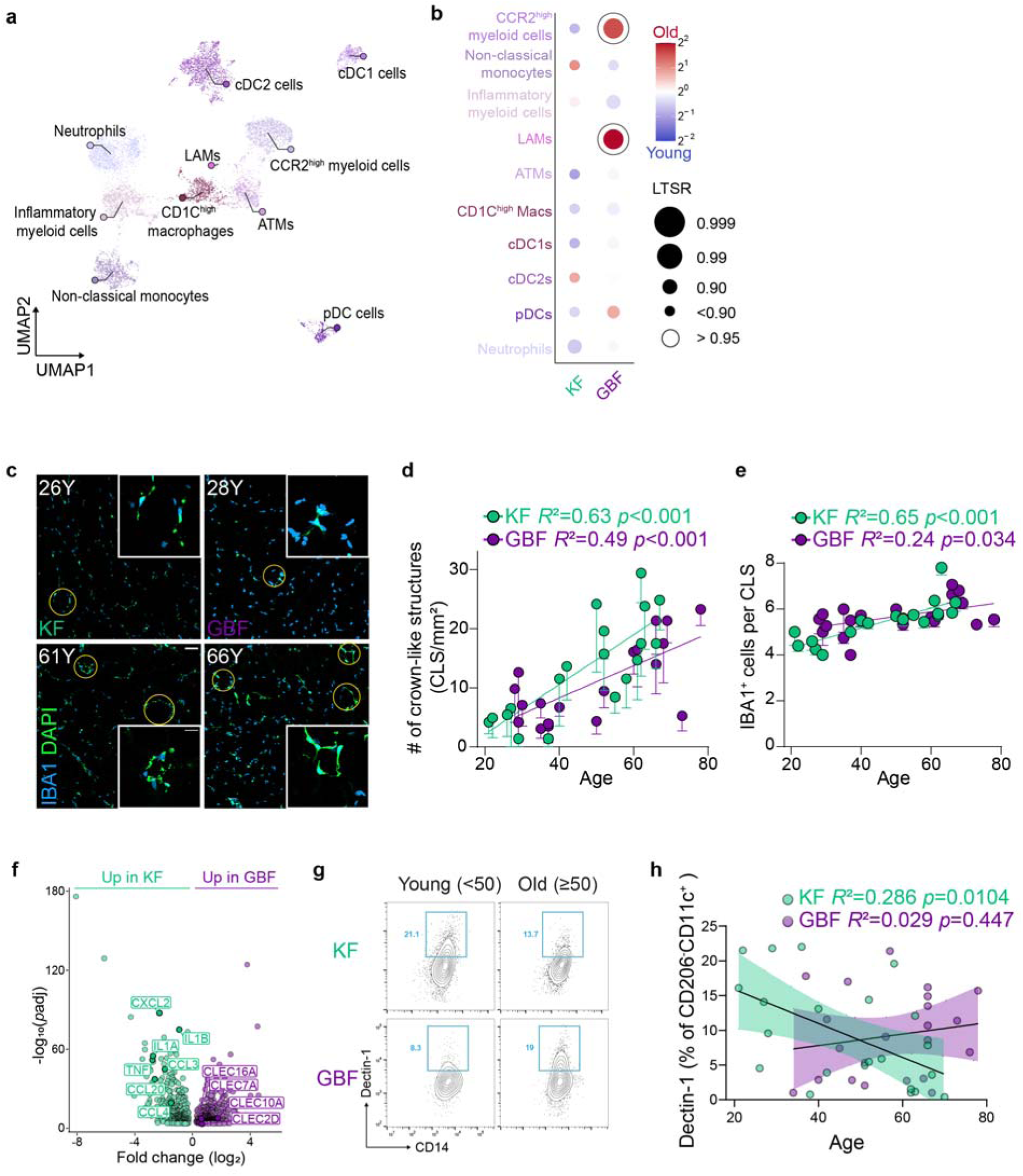
Distinct inflammatory program of VAT myeloid cells during aging. **a,** UMAP visualization of myeloid cells from the integrated VAT myeloid compartment, colored by subset. **b,** Bubble plot of the relative abundance of each myeloid subcluster in older versus younger donors, shown separately for the KF (first column) and GBF (second column). Color encodes log_2_ fold change between older and younger donors; bubble size encodes LTSR; open circles indicate LTSR > 0.95. **c,** Representative microscopic images of crown-like structures (CLS) in KF and GBF; donor age indicated. Scale bar = 50 µm; inset scale bar = 20 µm. **d,** CLS density as a function of donor age in KF and GBF. **e,** Number of IBA1^+^ cells per CLS versus donor age in the KF (green) and GBF (purple). **f,** Volcano plot of differentially expressed genes between GBF and KF inflammatory myeloid cells. **g,** Representative flow cytometry gating of Dectin-1^+^ cells among CD206^-^CD11c^+^ inflammatory myeloid cells across depots and ages. **h,** Dectin-1^+^ cell frequency (% of CD206^-^CD11c^+^) versus donor age in the KF (green) and GBF (purple). Shaded bands indicate 95% confidence intervals. For **d**, **e**, and **h**, *R*² and *p* value are indicated; lines, linear regression.

We next focused on inflammatory myeloid cells. During metabolic inflammation, recruited macrophages encircle dying adipocytes to form crown-like structures (CLS)^27,28^. We therefore examined whether age-associated myeloid accumulation in human VAT adopts this hallmark spatial organization of adipose inflammation. Immunohistochemical quantification revealed that CLS became more frequent and complex with age. The number of CLS per tissue area increased (**Fig. 5c,d**), and individual structures were composed of more myeloid cells, as IBA1^+^ counts per CLS also rose with age in both depots (**Fig. 5e**). Next, we asked whether their age-associated inflammatory program differs between depots. Thus, we performed differential gene expression analysis of the inflammatory myeloid cluster between the two depots. KF inflammatory myeloid cells showed higher expression of classical inflammatory genes including *IL1A*, *IL1B*, and *TNF*. In contrast, C-type lectin receptors (CLRs) including *CLEC7A*, *CLEC10A*, and *CLEC2D* were upregulated in GBF inflammatory myeloid cluster (**Fig. 5f**, **Extended Data Fig. 5h,i, Supplementary Table 6** for the full DEG list, and **Supplementary Table 7** for the full Metascape^29^ analysis result). Myeloid C-type lectin receptors recognize an array of ligands, from damage-associated molecular patterns to markers of altered self^30^. Among these, we focused on *CLEC7A* (Dectin-1), one of the best characterized CLRs and a sensor of endogenous ligands released or exposed upon cell injury^31–33^, for orthogonal validation. Consistent with differential gene expression analysis, flow cytometric analysis revealed that the frequency of Dectin-1^+^ in CD11c^+^ myeloid cells was decreased in the KF, while not changed in the GBF during aging (**Fig. 5g,h**). These changes in the frequency of Dectin-1^+^ myeloid cells during aging are conserved in a different type of myeloid cells (**Extended Data Fig. 5j**). Taken together, these results indicate that age-associated myeloid inflammation in human VAT is spatially organized and governed by a depot-specific molecular program.

### T cell-centric inflammaging as a hallmark of VAT immune remodeling

Our previous cell-type level analyses identified depot-specific shifts in inflammatory programs across multiple immune lineages with age. However, because human cohorts inevitably vary in BMI, sex, and other clinical covariates, we sought to determine which of these shifts reflect aging itself rather than associated confounders. To this end, we performed multivariate analyses of CD45^+^ immune cells comprising T cells, B cells, NKT cells, NK cells, macrophages, and dendritic cells characterized by flow cytometry across KF and GBF depots. Given the expected collinearity among immune cell lineages, we employed single-component partial least squares regression (PLSR), with age, immune lineages, BMI, and sex residualized prior to model fitting (see Methods). In the KF depot, the PLSR model revealed that T cells were the primary contributor to age-associated immune remodeling, as reflected by their high factor loadings on the first latent component (**Fig. 6a,b**). In the GBF depot, overall immune remodeling was less pronounced and did not attain statistical significance (**Fig. 6a**). To further validate and extend these findings, we applied a dimensionality reduction strategy using lineage-specific principal component analysis (PCA). For each immune block (macrophages, B cells, and T cells), the first principal component (PC1) was computed to capture the dominant axis of coordinated variation within that lineage (**Supplementary Table 8** and see Methods for more details). Partial Spearman correlation between each immune block PC1 and chronological age—adjusting for sex and BMI—confirmed that the T cell PC1 in the KF exhibited the strongest and most significant association with age (**Fig. 6c**). Although similar trends were observed in the GBF depot, these associations did not reach statistical significance (**Extended Data Fig. 6a**). Feature-level partial correlation analysis within the T cell compartment—encompassing αβ T (αβ T), MAIT, NKT, γδ1, and γδ2 T cell subsets—revealed that the proportion of αβ T cells was significantly and positively associated with aging (r = 0.673, q = 0.00576; **Fig. 6d**), whereas NKT cells, an innate-like T cell subset, displayed a significant negative association (r = −0.551, q = 0.0295; **Fig. 6e**). Partial correlation analyses incorporating additional clinical covariates further corroborated that T cell compositional shifts represent the most robust immune correlate of age in both depots relative to other immune lineages (**Extended Data Fig. 6b,c**). Collectively, these findings reveal a direct relationship between aging and VAT immune remodeling and highlight a T cell-centric shift as a hallmark of inflammaging in human VAT.

**Figure 6.**
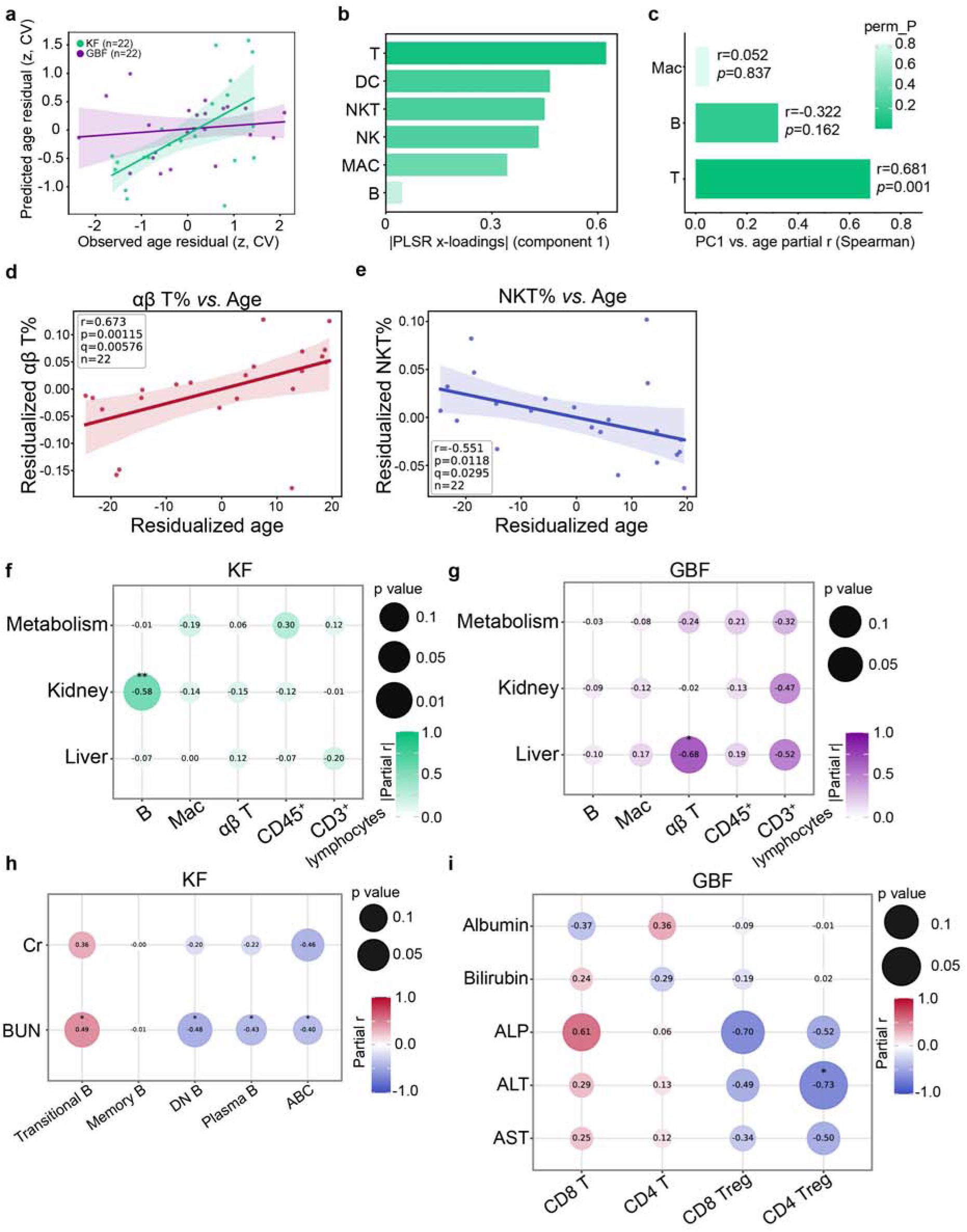
Depot-specific immune remodeling in human VAT with aging and functional coupling to adjacent organ status. **a**, Scatter plot of cross-validated predicted age residuals versus observed age residuals (z-scored) from the one-component PLSR model for the KF (green, n = 22) and GBF (purple, n = 22) depots. Shaded bands indicate 95% confidence intervals. All variables were pre-residualized for sex and BMI. **b**, Absolute value of PLSR first component x-loadings for each immune cell feature in the KF depot, ordered by absolute magnitude. **c**, Partial Spearman correlation coefficients between immune block PC1 and chronological age in the KF depot, adjusted for sex and BMI. Color intensity encodes permutation-derived *p*-values (perm_P). **d,e**, Partial Spearman correlation between residualized αβ T cell proportion (αβ T%;**d**) or NKT cell proportion (NKT%;**e**) and residualized age in the KF, adjusting for sex and BMI. Line, linear regression; shaded band, 95% confidence interval. **f,g**, Bubble plot of pairwise partial Spearman correlations between immune block PC1 and clinical block PC1 scores in the KF (**f**) and the GBF (**g**), adjusting for age and sex. Block-level associations are shown as absolute partial r values. Bubble size encodes *p*-value magnitude. Color encodes absolute value of partial r. **h**, Bubble plot of feature-level pairwise partial Spearman correlations between B cell subsets and kidney biomarkers in the KF depot, adjusting for sex and age. Bubble size encodes *p*-value. Color encodes partial r. **i**, Bubble plot of feature-level pairwise partial Spearman correlations between αβ T cell subsets and individual liver biomarkers in the GBF depot, adjusting for sex and age. * < 0.1; **q < 0.05.

### Depot-specific immune signatures reflect adjacent organ functional status

To determine whether depot-specific immune signatures reflect the functional status of anatomically adjacent organs, we performed partial Spearman correlation analyses between immune block PC1 scores and clinical block PC1 scores derived from blood biomarkers associated with liver, kidney, and metabolic functions. The kidney PC integrated markers of renal filtration capacity while the metabolism PC encompassed BMI and blood glucose/cholesterol. The liver PC integrated indicators of hepatocellular injury such as aspartate aminotransferase (AST) and alanine aminotransferase (ALT) (**Supplementary Table 8**). This block-level analysis revealed a spatially compartmentalized pattern of immune–organ interaction. In the KF depot, the B cell immune component was significantly correlated with the kidney clinical component (|r| = 0.58; **Fig. 6f**), whereas in the GBF depot, the αβ T cell component showed a strong association with the liver clinical component (|r| = 0.68; **Fig. 6g**). Feature-level pairwise analyses provided further mechanistic resolution. In the KF depot, the proportions of age-associated B cells (ABCs) and memory B cells were negatively correlated with BUN and creatinine levels, while transitional B cells exhibited positive correlations with these renal function markers (**Fig. 6h**). This association pattern was consistently stronger in the KF than in the GBF (**Extended Data Fig. 6d**). To further validate these observations, we examined partial correlations between B cell subsets and positive indicators of kidney function, including creatinine clearance (CrCl) and the estimated glomerular filtration rate (eGFR) calculated using the CKD-EPI equation^34^. Although these associations did not attain statistical significance, CrCl and eGFR displayed directionally consistent positive trends with activated B cell subsets (ABCs and plasma B cells) and negative trends with transitional B cells, mirroring the inverse relationships observed for BUN and creatinine (**Extended Data Fig. 6e**). Conversely, in the GBF depot, serum ALT levels were negatively correlated with CD4 regulatory T cells but showed a positive trend with non-regulatory αβ T cell subsets (**Fig. 6i**); these patterns were substantially weaker in the KF (**Extended Data Fig. 6f**). Taken together, these results demonstrate that the VAT immune landscape serves as a compartmentalized indicator of adjacent organ function, with depot-specific immune signatures partially reflecting the physiological status of the kidney and liver, respectively.

## Discussion

Here we define age-associated immune remodeling in human peri-organ VAT and show that VAT aging is anatomically compartmentalized across distinct fat depots. A central finding of this study is that aging in tissue-associated fat is not explained by uniform immune activation or by simple accumulation of terminally differentiated lymphocytes. Instead, aging was associated with a tissue-resident and granzyme K-associated inflammatory program in lymphocytes that varied between KF and GBF, and a spatially organized myeloid program that diverged between depots. These findings place VAT within the broader framework of human tissue immune aging while suggesting that metabolically active fat depots may impose distinct aging trajectories on resident and resident-like lymphocyte and myeloid compartments.

Recent studies of human tissue immunity have shown that tissue localization is a dominant determinant of immune cell identity and that T_RM_ cells can maintain site-specific phenotypes across age^19,35,36^. In those settings, circulating T_EM_ and T_EMRA_ cells acquire senescence-associated and *GZMK*-enriched signatures with age, whereas T_RM_ cells in mucosal and lymphoid tissues largely preserve resident identity without acquiring canonical immunosenescent features. Our data support the importance of tissue context but suggest that this apparent insulation of T_RM_ cells from inflammatory aging may be tissue-context dependent. In VAT, particularly in GBF, aging was associated with granzyme K induction across T_RM_-related, regulatory and innate-like lymphocyte compartments. At the transcriptomic level, *GZMK* marked selected CD8^+^ T_RM_ states, whereas flow cytometry revealed broader age-associated increases in granzyme K protein. This pattern suggests that granzyme K-associated remodeling in aging VAT reflects a tissue-level inflammatory adaptation, potentially shaped by adipose-specific cues such as lipid-rich metabolic stress and crown-like structure-associated myeloid inflammation, rather than a lineage-restricted or terminal cytotoxic senescence program. Consistent with this interpretation, granzyme K was preferentially associated with CD28-retaining cells, whereas granzyme B was enriched in CD28^-^populations. In GBF, enrichment of CD69^+^ memory Tregs, stronger granzyme K induction and expansion of an activated tissue-adapted Treg state further suggest qualitative remodeling of the regulatory compartment rather than simple Treg accumulation. Thus, granzyme K^+^ memory Tregs and related resident-like lymphocyte states may represent inflammation-adapted populations in which tissue adaptation, antigen experience and inflammatory responsiveness coexist. These findings also indicate that peri-organ VAT should not be treated as a single immunological compartment. GBF and KF followed distinct immune-aging trajectories, with GBF showing broader residency-associated lymphoid remodeling, higher CD103 expression and B cell enrichment, and KF showing greater myeloid representation, stronger age-associated B cell remodeling and compartmentalized changes within resident effector-like CD8^+^ T_RM_ states. This divergence suggests that local anatomical cues, including biliary/liver-adjacent or renal-adjacent stromal, vascular or metabolic signals, shape immune aging within the visceral fat compartment.

The age-associated increase in CD20^+^ T cells identifies a previously underappreciated axis of lymphocyte remodeling in VAT. CD20^+^ T cells are unlikely to be unique to VAT, as similar populations have been described in autoimmune and inflammatory settings^16,37,38^; however, their accumulation in aging VAT may reflect the distinctive features of this tissue niche, including local B cell enrichment, chronic antigenic or metabolic stimulation and close B-T cell interactions^39^. The absence of detectable CD20 transcripts in T cells, together with the acquisition of B cell-associated surface molecules, is consistent with membrane transfer through B-T cell contact rather than stable lineage-intrinsic CD20 expression. Notably, these cells showed a CD28-retaining, granzyme K-associated phenotype, suggesting that CD20 acquisition marks an antigen-experienced inflammatory memory state rather than terminal cytotoxic senescence. Future spatial and functional studies will be needed to determine whether CD20^+^ T cells preferentially accumulate in VAT relative to other tissues, whether they represent a consequence of the aging inflammatory niche or actively contribute to VAT inflammaging, and whether they participate in local immune regulation or inflammation during aging.

Although T cells emerged as a dominant correlate of VAT immune aging, myeloid cells remain central to the interpretation of adipose tissue inflammation. Relative myeloid frequencies in dissociated-cell datasets can be affected by lymphoid expansion, differential cell recovery and enzymatic loss of tissue-adherent populations. Consistent with this concern, *in situ* IBA1 staining and crown-like structure analysis revealed age-associated myeloid accumulation and spatial organization despite apparent relative myeloid decline in scRNA-seq and flow cytometric analyses. Moreover, myeloid remodeling was depot-specific: KF inflammatory myeloid cells showed a more classical *IL1A*/*IL1B*/*TNF*-high inflammatory profile, whereas GBF myeloid cells were enriched for C-type lectin receptor-associated genes, including *CLEC7A*. Thus, age-associated myeloid remodeling in VAT is both spatially organized and molecularly distinct between depots.

The association between VAT immune features and adjacent organ functional indices suggests a broader physiological role for peri-organ fat. In KF, B cell-related immune variation tracked with renal function-associated biomarkers, whereas in GBF, αβ T cell features were linked to liver-related clinical markers. These depot-specific associations raise the possibility that peri-organ VAT acts as a local immune sensor of adjacent organ status, rather than merely reflecting systemic inflammation, and are consistent with the broader need for site-specific immune monitoring in human disease and aging. Because the current study is cross-sectional, however, this model remains hypothesis-generating; longitudinal or perturbational studies will be needed to determine whether VAT immune remodeling contributes to organ dysfunction, responds to organ-derived cues or reflects bidirectional crosstalk between fat and adjacent tissues.

Together, our findings suggest that aging of human VAT is driven by depot-specific immune remodeling involving tissue-resident and resident-like lymphocytes, B-T cell interaction-associated states and spatially organized myeloid inflammation. Rather than representing a uniform macrophage-centered inflammatory process, VAT aging involves a T cell-centric and granzyme K-associated program that is shaped by local adipose tissue niches and adjacent organ context. These observations highlight the need to resolve adipose tissue immunity by anatomical depot and suggest that peri-organ VAT may provide a window into local organ-associated immune aging.

## Materials and Methods

### Human adipose tissue specimens and clinical data

Human gallbladder (GB)-associated visceral adipose tissue (VAT) was obtained from asymptomatic patients undergoing elective laparoscopic cholecystectomy for incidental gallbladder polyps at Korea University Anam Hospital. Eligibility was restricted to patients without clinical or radiographic evidence of intra-abdominal infection or inflammation. Patients with acute cholecystitis were excluded at the enrollment stage, and those with histopathologically confirmed gallbladder malignancy on the resected specimen were subsequently excluded. Human kidney-associated VAT samples were obtained from healthy living donors undergoing donor nephrectomy for kidney transplantation at Asan Medical Center, Seoul. Donors with radiographic evidence of intra-abdominal inflammation also were excluded. Donor inclusion was conducted under the following criteria to minimize confounding from systemic disease and immunomodulatory exposures. All donors were non-obese (18 < BMI < 30), had no prior abdominal surgery, and had no history of cardiovascular disease, chronic liver disease, chronic kidney disease, active malignancy, or systemic immunological disorders, including autoimmune or chronic inflammatory conditions. Donors receiving medications known to alter immune function including systemic corticosteroids, immunosuppressants, and chronic non-steroidal anti-inflammatory drugs were excluded. For common metabolic comorbidities (hypertension, type 2 diabetes mellitus, and dyslipidemia), only well-controlled cases on non-immunomodulatory therapy were included. Detailed donor characteristics, including age, sex, BMI, comorbidities, medications, and laboratory values, are provided in Supplementary Table 1. Routine clinical laboratory parameters measuring metabolic, hepatic, and renal function were obtained at baseline as part of standard preoperative evaluation; values used in downstream analyses are listed in Supplementary Table 8.

The study was approved by the Institutional Review Boards of Korea University Anam Hospital (IRB No. 2022AN0294) and Asan Medical Center (IRB No. 2024-0881) and was conducted in accordance with the Declaration of Helsinki. Written informed consent was obtained from all donors prior to sample collection.

### Stromal vascular fraction (SVF) isolation and sample handling

SVF cells were isolated from GBF and KF samples by collagenase-mediated digestion using tissue-processing workflows adapted from a published protocol^40^. Briefly, adipose tissue was washed in cold washing buffer consisting of DPBS containing 2% FBS and minced in chilled washing buffer. For both depots, freshly minced tissue was digested with collagenase I (2,500 U/g tissue) and DNase I (20 U/g tissue) at 37 °C with agitation at 250 rpm for 60 min. Digested suspensions were filtered through 100 µm and 70 µm cell strainers and centrifuged at 800 × g for 10 min at 4 °C to recover the SVF fraction. Red blood cells were lysed using RBC lysis buffer (BioLegend, 420301), and cells were washed and resuspended in washing buffer before downstream processing. For single-cell RNA-seq, GBF-derived SVF cells were processed freshly after digestion. KF-derived SVF cells were cryopreserved as SVF cell suspensions after fresh tissue digestion in Recovery Cell Culture Freezing Medium using a Mr. Frosty freezing container at −80 °C, and were later thawed for downstream single-cell processing. GBF- and KF-derived SVF cells were washed in DPBS containing 2% FBS and incubated with Fc receptor blocking reagent (BioLegend, 422301). For KF-derived SVF samples, cell hashing was performed using TotalSeq-C anti-human hashtag antibodies (BioLegend). Both GBF- and KF-derived SVF cells were stained with CD45-PE antibody (BioLegend, 304058) and DAPI (Sigma-Aldrich, D9542). Viable CD45^+^ immune cells were isolated by fluorescence-activated cell sorting using a MA900 Multi-Application Cell Sorter (Sony, Japan). After sorting, hashtagged KF samples were pooled at four samples per 10x Genomics run across two independent runs. GBF-derived SVF samples were processed individually without hashing.

### Single-cell RNA-sequencing library preparation and sequencing

Sorted viable CD45^+^ immune cells were loaded onto the 10x Genomics Chromium platform for GEM generation and barcoding according to the manufacturer’s instructions. GBF-derived SVF samples were processed using Chromium Next GEM Single Cell 3′ v3.1 chemistry, whereas KF-derived SVF samples were processed using Chromium GEM-X Single Cell 5′ v3 chemistry. Library quality and fragment size distribution were assessed using an Agilent Bioanalyzer DNA chip before sequencing. GBF-derived SVF libraries were sequenced using HiSeq X Ten or Illumina NovaSeq 6000 platforms. KF-derived SVF libraries were sequenced using the Illumina NextSeq 2000 platform.

Raw sequencing data were processed using Cell Ranger. GBF-derived SVF libraries were processed individually using the standard Cell Ranger count workflow v7.1.0, whereas KF-derived SVF libraries were processed using Cell Ranger multi v9.0.1 to jointly process gene expression and TotalSeq-C hashtag libraries. Reads were aligned to the human GRCh38 reference genome (refdata-gex-GRCh38-2020-A). For KF-derived multiplexed samples, Cell Ranger multi sample assignments were used for demultiplexing.

Cell Ranger output matrices were corrected for ambient RNA contamination using SoupX v1.6.2^41^. Corrected matrices were analyzed in R using Seurat v5.0.1^42^. Before Seurat quality-control filtering, 148,353 cell barcodes were included. Seurat objects were generated using genes detected in at least three cells and cells with at least 200 detected features. Cells were further filtered based on RNA counts, detected features, mitochondrial gene content and ribosomal gene content. The final filtering thresholds were 300–35,000 RNA counts per cell, 200–7,000 detected RNA features per cell, <10% mitochondrial gene content and <50% ribosomal gene content. Mitochondrial gene content was calculated using genes beginning with MT-, and ribosomal gene content was calculated from ribosomal protein genes. After quality-control filtering, 116,307 cells were retained.

Doublets were identified independently for each biological sample using DoubletFinder v2.0.6^43^ with an expected doublet rate of 7.5% and homotypic adjustment. After doublet removal, 109,465 singlets were retained for integration.

The remaining singlet cells were normalized using log normalization with a scale factor of 10,000. Highly variable genes were identified using the variance-stabilizing transformation method, selecting 3,000 variable features. Mitochondrial genes, ribosomal genes, sex-linked genes, hemoglobin genes and MALAT1 were excluded from the variable feature set before dimensionality reduction. Data were scaled with regression of mitochondrial gene percentage and number of detected features before principal component analysis.

Singlet cells from all GBF- and KF-derived SVF samples were integrated using Harmony v1.2.3^13^ with sample identity as the integration variable. The first 50 Harmony dimensions were used for UMAP visualization, graph construction and graph-based clustering in Seurat. Clusters were identified at resolution 0.3.

### Cell-type annotation and downstream single-cell analyses

Cell clusters were annotated manually based on canonical immune lineage markers and inspection of marker expression across the integrated UMAP. Cluster-enriched genes were identified using Seurat’s FindAllMarkers function by comparing each annotated cluster with all other cells. Only positive markers were considered, with a minimum expression fraction of 25% and a log2 fold-change threshold of 0.5. Significant markers were defined using an adjusted P value <0.05, and the top 100 significant cluster-enriched genes ranked by average log2 fold change were used to guide cell-type annotation together with canonical immune-lineage markers, dot plots and feature plots.

Major immune populations were identified using markers including *PTPRC*, *CD3E*, *CD4*, *CD8A*, *CD79A*, *CD68*, *CD14*, *LYZ*, *FCGR3A* and *NCAM1*. Potential non-immune contaminating clusters were assessed using stromal, endothelial and adipocyte-associated markers, including *DCN*, *COL1A1*, *VWF*, *FLT1*, *EMCN* and *FABP4*. After exclusion of non-immune contaminating clusters, 96,341 immune cells were retained for downstream analyses.

For lineage-level analyses, annotated immune populations were subsetted and reprocessed independently. Subsetted objects were log-normalized, variable features were identified using the variance-stabilizing transformation method, and data were scaled before principal component analysis. Mitochondrial gene percentage and UMI count were regressed during scaling. Subsetted objects were integrated using Harmony with sample identity as the integration variable, and UMAP visualization, graph construction and clustering were performed using the first 50 Harmony dimensions. Subclusters were annotated using lineage-specific canonical markers, cluster-enriched genes, dot plots and feature plots.

### Cell-type composition analysis

Cell-type composition was analyzed using donor/sample-level counts derived from annotated cell identities. For each analyzed cell set, cells were aggregated by sample, fat depot, age, and annotated identity. Descriptive frequencies were calculated as the proportion of cells assigned to each annotated identity within the corresponding analyzed cell set, and descriptive fold changes were calculated between old and young donors within each fat depot or between KF and GBF where indicated. To reduce instability from rare populations, descriptive fold-change estimates required detection in at least two non-zero donors in either comparison group and a minimum contribution of 0.1% to the analyzed cell set. Fold changes involving zero frequencies in one comparison group were capped at ±2 log2 fold change for visualization. Statistical support for composition changes was estimated using a count-based mixed-model framework adapted from previously described single-cell composition analyses. Cell-type-specific counts were modeled using negative-binomial generalized linear mixed models with a log link. Donor/sample identity was included as a random intercept, and an offset for the log-transformed total number of cells recovered from the corresponding sample within the analyzed cell set was included to model relative abundance.

Age-associated changes were modeled separately within GBF and KF. Age was scaled and centered, and cell-type-specific age effects were estimated using a CellType × Age interaction: Ncs ∼ 0 + CellType + CellType:Age_scaled + offset(log(total_cells_sample)) + (1 | Sample), where Ncs denotes cell-type-specific counts and Age_scaled denotes scaled and centered donor age. Depot-associated differences between KF and GBF were modeled analogously using: Ncs ∼ 0 + CellType + CellType:Depot + offset(log(total_cells_sample)) + (1 | Sample). Because depot was partially confounded with tissue processing, 10x chemistry, and sequencing platform, depot-associated effects were interpreted descriptively as depot-associated abundance differences. For each annotated identity, local true sign rate (LTSR) was estimated from the model-derived coefficient and standard error as pnorm(abs(beta / SE)), representing a normal-approximation probability that the inferred direction of effect was supported by the fitted model. LTSR values range from 0 to 1, with higher values indicating stronger directional support; changes with LTSR ≥ 0.95 were considered strongly supported.

Pairwise differential expression analyses, including lineage- or subset-specific comparisons in myeloid and B-cell populations, were performed using Seurat’s FindMarkers function on normalized gene expression values. Logistic regression was used with latent-variable adjustment for sex, mitochondrial gene percentage, ribosomal gene percentage and UMI count. Genes with an absolute log2 fold change of at least 0.25 and an adjusted P value <0.01 were considered significant. For each pairwise comparison, the direction of average log2 fold change was interpreted according to the order of the groups specified in Seurat, with positive values indicating enrichment in the tested group and negative values indicating enrichment in the reference group.

### Functional enrichment analysis

Functional enrichment analysis was performed using Metascape, a web-based platform for gene-list annotation and pathway enrichment analysis^29^. For each pairwise comparison, genes passing the differential expression threshold of adjusted P value <0.01 were ranked by log2 fold change. The top 100 upregulated and top 100 downregulated genes were analyzed separately in Metascape. Enriched biological terms and pathway groups were exported from Metascape and used for downstream visualization and interpretation.

### Immunohistochemistry

Adipose tissue samples were washed three times in cold 2% FBS in DPBS and fixed in 4% paraformaldehyde at 4 °C for 48 h. Following fixation, samples were washed in DPBS and cryoprotected in 30% sucrose in PBS for 48 h. Tissues were embedded in frozen-section embedding medium compound and stored at −80 °C until sectioning. Frozen sections (25–40 µm) were generated using a cryostat at −21 to −24 °C, mounted onto slides, and stored at −20 °C or −80 °C until staining. Sections were thawed at room temperature, blocked for 1 h in staining buffer consisting of PBS containing 5% normal goat serum and 0.1% Triton X-100, and incubated overnight at 4 °C with anti-IBA1 primary antibody (234-308, SYSY, 1:500) diluted in the same staining buffer. After washing, sections were incubated with Goat anti-Guinea Pig IgG (H+L) Highly Cross-Adsorbed Secondary Antibody conjugated to Alexa Fluor 488 for IBA1 detection (Invitrogen/Thermo Fisher Scientific, A-11073), diluted 1:500 in staining buffer for 1 h at room temperature. Nuclei were counterstained with DAPI using Fluoromount-G (SouthernBiotech, 0100-20). Images were acquired using LSM880 (Carl Zeiss, Germany) with identical acquisition settings across samples. Details for all reagents used for immunofluorescence staining are provided in Supplementary Table 9.

### Image analysis and quantification

Quantification of immunofluorescence signals was performed using custom macros in ImageJ (NIH). Briefly, confocal images were imported and split into individual channels, followed by maximum-intensity projection where applicable. Images were converted to 8-bit and processed using consistent intensity thresholds across all samples. Cell segmentation was performed using automated thresholding methods optimized for each channel, including Yen thresholding for DAPI and MaxEntropy thresholding for IBA1, combined with background subtraction, median filtering and morphological operations, including opening and watershed, to refine object detection. IBA1^+^ signal and DAPI^+^ nuclei were segmented using consistent thresholding parameters, and regions of interest (ROIs) were quantified for cell number, area and signal intensity where applicable.

To validate automated quantification, a subset of images was independently assessed in a blinded manner by two experimenters using manual counting. All measurements were performed using identical parameters across samples. Quality-control images with ROI overlays were generated to validate segmentation accuracy. For each biological sample, 6–10 images were analyzed for IBA1 and DAPI quantification.

Crown-like structures (CLSs) were defined as adipocytes surrounded by aggregates of ≥4 IBA1^+^/DAPI^+^ cells. CLS abundance was quantified by counting the number of CLSs in each field of view and averaging across 6–10 fields of view to obtain the mean CLS count per field of view for each biological sample. For CLS composition analysis, DAPI^+^ nuclei and IBA1^+^ cells within CLS regions were manually counted.

### TMT-based quantitative proteomics by mass spectrometry

Fresh peri-gallbladder visceral adipose tissue was processed using the same stromal vascular fraction isolation workflow used for single-cell RNA-sequencing samples. Briefly, freshly collected adipose tissue was mechanically minced and enzymatically digested with collagenase I at 37°C with agitation. The resulting cell suspension was filtered, washed and resuspended for fluorescence-activated cell sorting. Cells were incubated with Fc receptor blocking reagent to reduce nonspecific antibody binding and then stained with Zombie NIR viability dye to exclude dead cells, followed by surface staining with antibodies against CD45, CD3, CD4, CD8, CD62L, CD45RO, CD25, CD127 and CD56. Viable CD45^+^ lymphocytes were identified by forward- and side-scatter properties and exclusion of Zombie NIR^+^ events. T cells were gated as CD45^+^CD3^+^ events, and CD4 and CD8 T-cell fractions were subsequently identified. Effector/memory-enriched CD8 T cells were sorted as viable CD45^+^CD3^+^CD8^+^CD4^-^CD45RO^+^CD62L^-^ cells. Conventional non-regulatory CD4 T cells were sorted as viable CD45^+^CD3^+^CD4^+^CD8^-^CD25^-^CD127^+^ cells. CD56^+^ events were excluded where applicable to minimize contamination by NK and NKT-like cells. Sorted populations were collected for downstream proteomic analysis using MA900 Multi-Application Cell Sorter (Sony, Japan).

For protein extraction, the sorted cells in lysis buffer (2% SDS, 2 mM MgCl_2_, 50 mM triethylammonium bicarbonate (TEAB), 1X Halt protease and phosphatase inhibitor cocktail) were disrupted using a Bioruptor Sonicator (Diagenode, Denvile, NJ). Then, 0.5 µL of 250 units/µL benzonase nuclease was added to the cell lysates. The mixture was heated at 95 °C for 5 minutes and cooled to room temperature (RT). Samples were incubated with 20 mM dithiothreitol (DTT) at 60°C for 10 minutes to reduce disulfide bonds, followed by alkylation with 40 mM iodoacetamide at RT for 30 minutes in the dark. The alkylation reaction was quenched by adding additional DTT to a final concentration of 10 mM. Subsequently, on-column enzymatic digestion was carried out using an S-Trap column according to the manufacturer’s protocol (PROTIFI, Fairport, NY), using trypsin/Lys-C mix (Promega, Madison, WI) at 47°C for 1 h. The resultant peptides were labeled with 17-plex TMT isobaric reagents (ThermoFisher Scientific, Waltham, MA) as described previously^44^. The labeled samples were then combined and dried using a vacuum concentrator. The combined peptide sample was reconstituted in 0.1% formic acid and subjected to LC-MS/MS analysis using a Q-Exactive mass spectrometer coupled to an UltiMate 3000 HPLC system (Thermo Fisher Scientific). Peptides were separated on an EASY-Spray analytical column (50 cm × 75 μm inner diameter, C18, 2 μm particle size, Thermo Fisher Scientific). The mobile phase consisted of solvent A (0.1% formic acid in water) and solvent B (0.1% formic acid in 80% acetonitrile). Peptide separation was achieved using a linear gradient from 2% to 40% solvent B over 140 minutes, at a flow rate of 300 nL/min. Instrument settings were consistent with those reported previously^44^ for 50-cm column usage, except for the following adjustments: MS1 scan range of 300–1800 m/z and a spray voltage of 2.5 kV.

The raw MS data files were initially converted to mzML format using MSConvert (ProteoWizard). Subsequent isobaric quantification based on TMT labeling was carried out utilizing MSFragger (v4.1) and IonQuant (v1.10.27) within the FragPipe environment (v22.0), using the UniProt Homo sapiens reference proteome (release 2024_01) for database searching. The built-in workflow “TMT16” was applied with customized settings: within the Quant (Isobaric) module, the min purity threshold was set to 0.75, and the label type was set to TMT-18.

### Bioinformatics analysis of proteomic data

Bioinformatic analyses were performed using Perseus (v1.6.14.0) in combination with R (v4.3.2) executed within the R-Studio environment (v2024.12.0). The raw quantitative data file was imported into Perseus for downstream preprocessing. To account for differences in sample input amounts, protein-level TMT reporter ion intensities were normalized to the total reporter ion intensity of each sample by dividing individual protein intensities by the corresponding sample-wise intensity sum. The normalized values were subsequently log2-transformed. Within each CD8 and CD4 cell population, statistical comparisons between old and young groups were performed using a two-sided Student’s t-test in Perseus. The resulting datasets, including protein intensities, fold changes, and p-values, were then exported for additional downstream analyses conducted in R.

For quality control (QC), data visualizations including intensity density plots and boxplots were generated in R using ggplot2, while correlation scatter plots were produced using ggpubr. Subcellular localization boxplots were generated by first classifying proteins according to their annotated cellular localization using the DAVID platform (https://david.ncifcrf.gov/, accessed on 30 April 2025), followed by visualization in R using ggplot2.

Pathway analyses were performed using gene sets from the MSigDB C5 Biological Process (BP) collection. Gene set analysis was conducted in R using the *runGSA* from the *piano* package, with pathway activity scores calculated using the PAGE method. For downstream interpretation, analyses were restricted to mitochondria-related pathways by selecting gene sets annotated with mitochondrial-associated terms, including “MITOCHONDR”, “OXIDATIVE”, “RESPIRATORY”, “TCA”, “ATP”, “ELECTRON_TRANSPORT”, and “MITO”.

To visualize the expression patterns of proteins associated with CD8 T cell effector function, a predefined gene list corresponding to “cell killing” biological processes was retrieved from the MSigDB C5 BP database. Protein expression values were sorted based on log2 fold change (old/young), and the resulting data were visualized as a heatmap using the *pheatmap* package in R.

### Flow cytometry analysis

For flow cytometry experiments, SVF cells from both depots were first isolated by collagenase-mediated digestion as described above, cryopreserved as cell suspensions in Recovery Cell Culture Freezing Medium using a Mr. Frosty freezing container at −80 °C, and subsequently transferred to liquid nitrogen for long-term storage. SVF preservation conditions were optimized through validation experiments comparing fresh and cryopreserved SVF-processing workflows to maintain immune-cell representation. For flow cytometry staining, cryopreserved SVF cell suspensions were washed twice and rested in complete medium consisting of RPMI medium supplemented with 10% FBS, 20 mM HEPES and 1% penicillin/streptomycin for 2 h at 37 °C before staining. For live-dead staining, cells were incubated with Zombie UV Fixable Viability Kit (Biolegend, 423107) in PBS for 20 min at 4°C. Then, the cells were washed with FACS buffer (PBS containing 1% BSA and 0.05% Sodium Azide) and incubated with Human BD Fc block (BD biosciences, 564220) for 10 min at 4°C. For surface staining, cells were incubated with surface antibodies in FACS buffer for 30 min at 4°C. For intracellular staining, cells were then fixed and permeabilized using Foxp3/Transcription Factor Staining Buffer Kit (Tonbo, TNB-0607-KIT) for detection of Foxp3 and BD Cytofix/Cytoperm Fixation/Permeabilization Kit (BD biosciences, 554714) for detection of Granzyme B, Granzyme K and Ki-67 according to the manufacturer’s instructions for 45 min at 4°C, followed by overnight incubation with intracellular antibodies in the corresponding Perm/Wash buffer at 4°C. The detailed antibody information is documented in Supplementary Table 9. Unbound antibodies were removed by washing, and cells were resuspended in FACS buffer containing Precision Count Beads (Biolegend, 424902) for absolute cell counting. Flow cytometry was performed using CytoFLEX LX flow cytometer (Beckman Coulter). Data were analyzed using FlowJo software v10.4 and t-SNE plots were generated using OMIQ with the following parameters: perplexity = 30, num result components = 2, iterations = 1000 and learning rate = 5000.

## Data availability

The mass spectrometry proteomics data have been deposited to the ProteomeXchange Consortium via the PRIDE partner repository with the dataset identifier PXD073404 (doi.org/10.6019/PXD073404). The data are also available under the accession number KAP242385 at the Korea BioData Station (K-BDS, https://kbds.re.kr).

## Acknowledgments

We are grateful to Byungjin Hwang (Yonsei University) for help with scRNA-seq multiplexing methodology as well as Yuna Chun (Chayeon Science) for help with the FACS sorting. We are grateful to the R.T. Han and Y.P. labs for helpful comments on the manuscript.

We acknowledge the following financial support for the research, and/or publication of this Article. R.T.H. was supported by the Ministry of Health & Welfare and Ministry of Science and ICT, Republic of Korea (RS-2024-00346245, RS-2025-25458875, RS-2025-0230482, RS-2025-25462906, RS-2024-00449882, and RS-2026-25523723). R.C. was supported by the Ministry of Health & Welfare, Republic of Korea (RS-2025-24534526). H.J.L. was supported by the Ministry of Science and ICT, Republic of Korea (RS-2025-02305395). Y.P. was supported by the KIST Institutional Program and by grants from the National Research Foundation of Korea (RS-2024-00337093 and RS-2026-25504579). H.S.J. was supported by the Ministry of Health & Welfare, Republic of Korea (RS-2025-25460074). H.R. was supported by the National Research Foundation of Korea (NRF) (RS-2024-00405650 and RS-2026-25487445).

## Author information

These authors contributed equally: Rodrigo Castaneda, Hye-In Sim.

These authors jointly supervised this work: Youngmin Ko, Hye-Sung Jo, Yoon Park, and Rafael T. Han.

## Contributions

R.T.H. and Y.P. conceived the study and supervised the work. Y.K. and H.S.J. acquired organ-donor tissue and designed the study cohort; K.J.P. assisted Y.K. with cohort design and analyzed patient radiological examinations for exclusion criteria. R.C. preserved collected tissues, performed scRNA-seq tissue processing and experimental workflows, carried out wet-lab experiments and interpreted scRNA-seq data. H.I.S. performed flow cytometry experiments and interpreted data. H.J.L. performed immunohistochemistry and manual image analysis. H.J.K. assisted R.C. with integrating the scRNA-seq data. H.Y.J. and H.R. assisted R.C. with scRNA-seq analysis of B cells and T cells, respectively. B.Y.J. assisted R.C. with multiplexed scRNA-seq analysis. N.P. and C.L. performed proteomics experiments and interpreted data. M.Y. and K.R. performed the clinical-biomarker correlation analyses. Interpretation of the results: C.L., K.R., H.R., H.Y.J, Y.P. and R.T.H. Manuscript writing: C.L., K.R., Y.K., H.S.J., H.K.S, Y.P. and R.T.H. All authors reviewed and approved the manuscript.

---

**Extended Data Figure 1.**
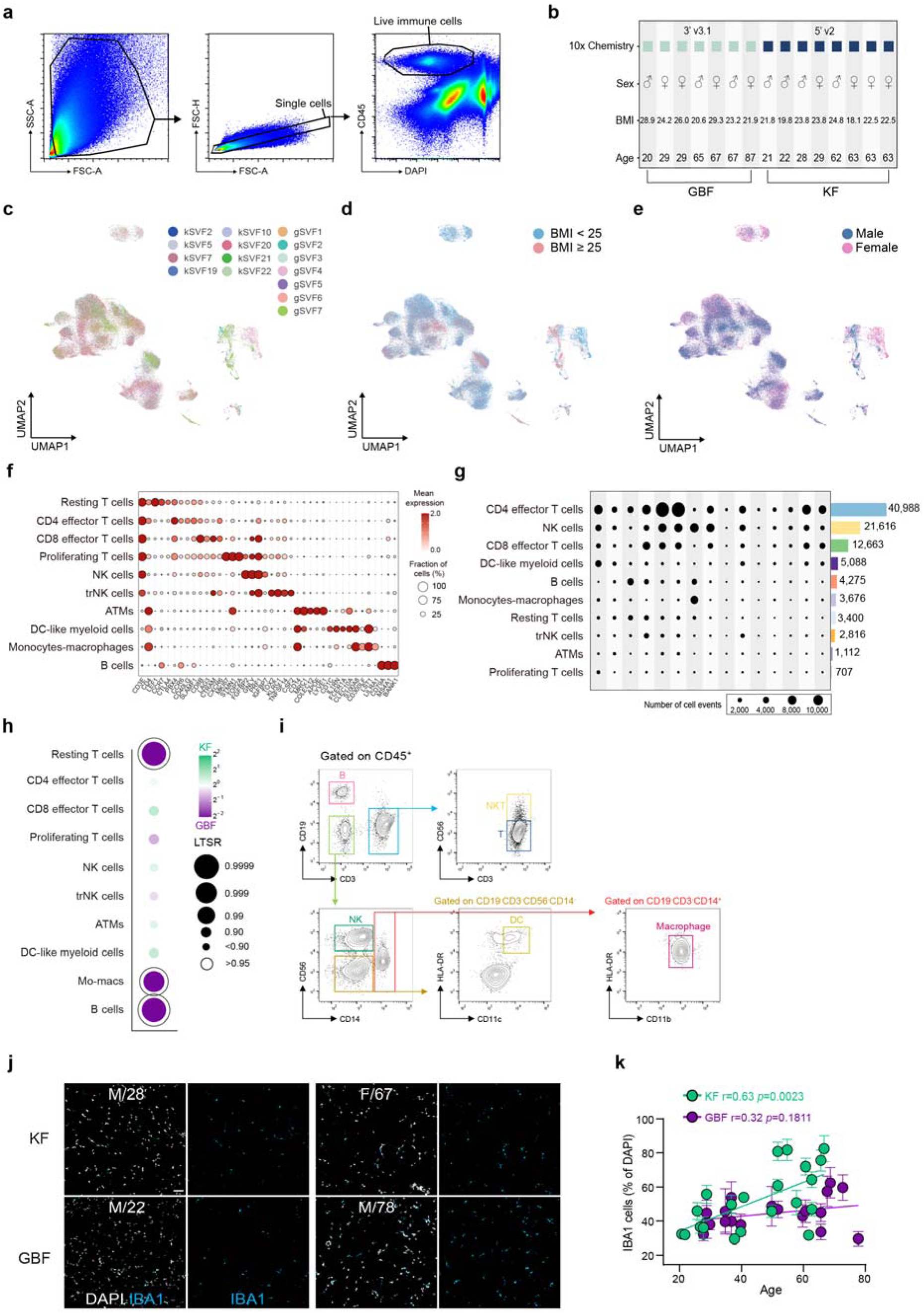
Quality control, donor metadata, and cell cluster characterization for the VAT immune atlas. a, Flow cytometry gating strategy for isolation of live CD45^+^ immune cells from VAT. **b,** Donor metadata for the scRNA-seq cohort, grouped by depot (GBF, left; KF, right). Rows indicate 10x Genomics library chemistry, sex, BMI, and age. **c-e,** UMAP colored by donor identity (**c**), BMI category (**d**; BMI < 25; BMI ≥ 25), and sex (**e**). **f,** Dot plot of canonical marker gene expression across the 10 annotated immune cell clusters. Dot size encodes fraction of expressing cells (%); color encodes mean expression. **g,** Dot plot of cell event numbers per donor per cluster; dot size encodes the number of cell events. Numbers at right indicate the total cell count per cluster. **h,** Bubble plot of depot-level enrichment for each cell cluster. Color encodes log_2_ fold change between KF and GBF. Bubble size encodes LTSR; open circles indicate LTSR > 0.95. **i**, Representative flow cytometry gating plots used to define major CD45^+^ immune cell subsets in VAT. **j,** Representative immunofluorescence images of IBA1^+^ myeloid cells and DAPI nuclei in KF and GBF (donor sex/age indicated). Scale bar = 50 μm. **k,** IBA1^+^ cell density (% of DAPI^+^ cells) versus donor age in the KF (green) and GBF (purple) depots. Each point is a donor (mean ± s.e.m. across imaged fields); Pearson *r* and *p*-values are shown.

**Extended Data Figure 2.**
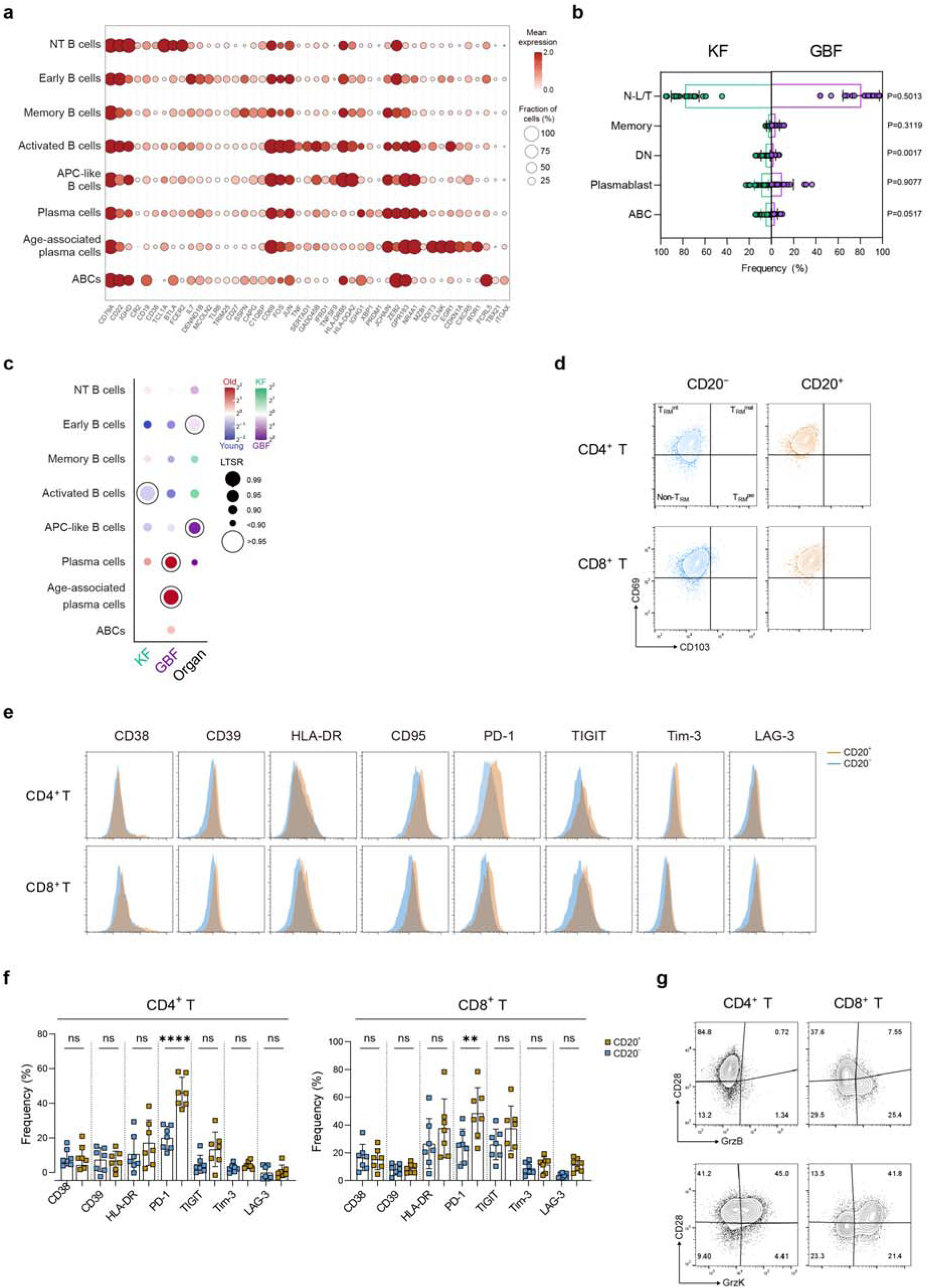
B cell heterogeneity and CD20^+^ T cell phenotypes in VAT. a, Dot plot of marker gene expression across annotated B-lineage subclusters. Dot size encodes the fraction of cells expressing each gene (%); color encodes mean scaled expression. **b**, Summary plots showing the frequencies of the indicated CD19^+^ B cell subsets in KF and GBF. **c**, Bubble plot of age- and depot-level differential abundance for each B cell subcluster. For the KF and GBF columns, color encodes log fold change between older and younger donors (red, old; blue, young); for the Organ column, color encodes log fold change between depots (green, KF; purple, GBF). Bubble size encodes the local true sign rate (LTSR), a measure of statistical confidence ranging from 0 to 1; open circles indicate LTSR > 0.95. **d**, Representative flow cytometry plots showing CD69/CD103-defined T_RM_-related states among CD20^+^ and CD20^-^CD4^+^ and CD8^+^ T cells. **e**, Representative histograms showing expression of CD38, CD39, HLA-DR, CD95, PD-1, TIGIT, TIM-3 and LAG-3 in CD20^+^ and CD20^-^ CD4^+^ and CD8^+^ T cells. **f**, Summary plots showing the frequencies of CD38^+^, CD39^+^, HLA-DR^+^, CD95^+^, PD-1^+^, TIGIT^+^, TIM-3^+^ and LAG-3^+^ cells among CD20^+^ and CD20^-^ CD4^+^ and CD8^+^ T cells in KF (n = 7 donors). **g**, Representative flow cytometry plots showing CD28 and granzyme B co-expression and CD28 and granzyme K co-expression in CD4^+^ and CD8^+^ T cells. Data in **b** and **f** are mean ± s.d. *p* value were determined by two-tailed unpaired *t*-test. \*\**p* < 0.01, \*\*\*\**p* < 0.0001; ns, not significant.

**Extended Data Figure 3.**
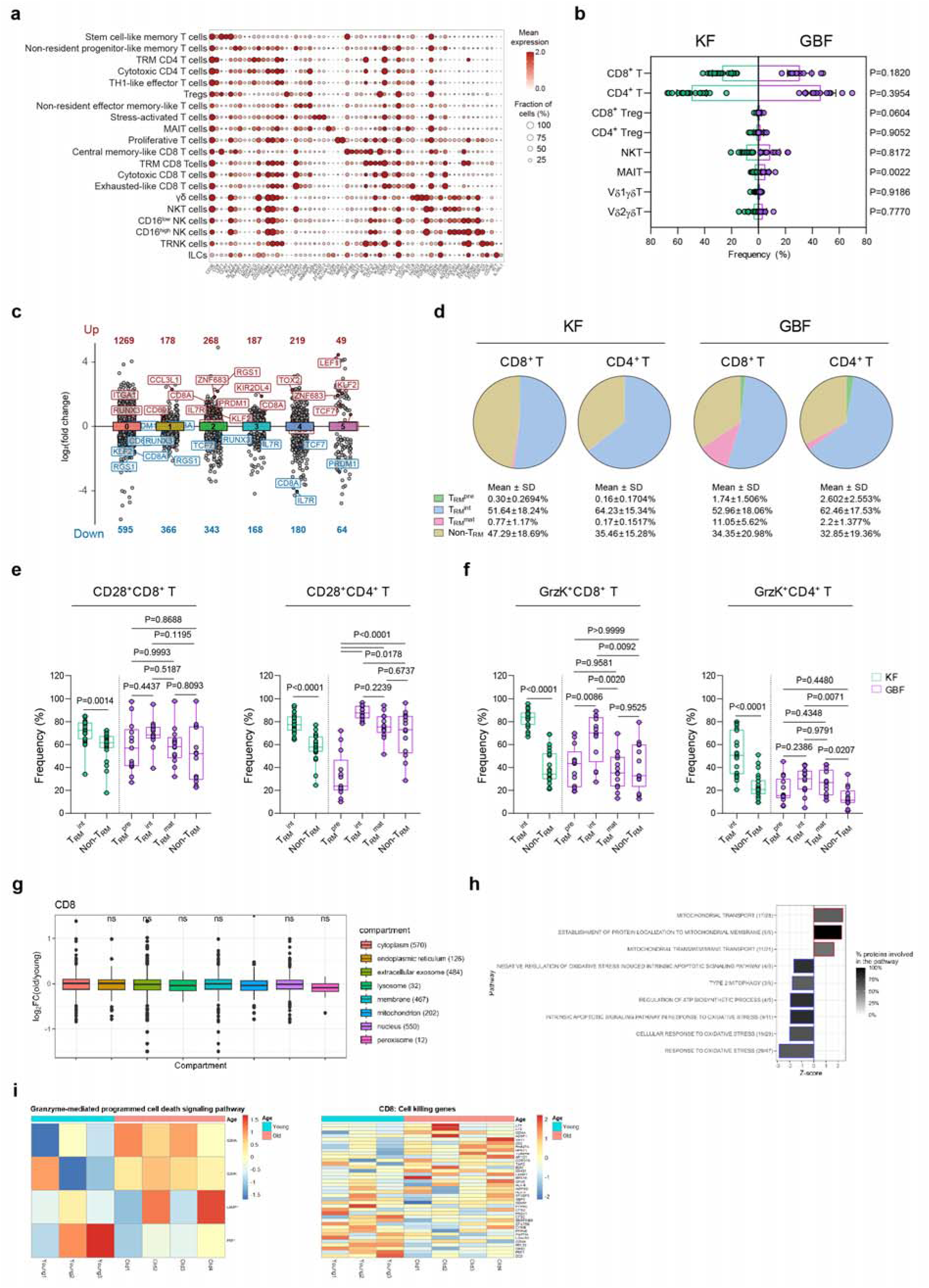
T cell subset heterogeneity and T_RM_-associated remodeling in VAT. **a**, Dot plot of selected marker genes across the 20 annotated T, NK, and innate lymphoid subclusters. Dot size indicates the fraction of cells expressing each gene, and color indicates mean expression. **b**, Box plots showing the frequencies of the indicated CD3^+^ T cell subsets in KF and GBF. **c**, Marker genes for the six T_RM_ subclusters (clusters 0–5, corresponding to T_RM_0–T_RM_5, each compared against all other cells). Points represent individual genes (adjusted *p* < 0.01), positioned by log fold change. The numbers above and below each cluster give the count of enriched (red) and depleted (blue) genes. **d**, Pie charts showing the mean proportional composition of CD69/CD103-defined T_RM_-related states among CD8^+^ and CD4^+^ T cells in KF and GBF. **e,f**, Box plots showing the frequencies of CD28^+^ cells (e) and granzyme K^+^ cells (f) within each CD69/CD103-defined T_RM_-related state among CD8^+^ and CD4^+^ T cells in KF and GBF. **g**, Box plots showing log fold changes in intracellular proteins between 3 younger and 4 older donors in CD8^+^ T cells from GBF, grouped by subcellular localization assigned using DAVID annotation. **h**, Pathway-level analysis of mitochondrial-associated biological processes in CD8^+^ T cells from GBF, based on PAGE. **i**, Heatmaps showing protein expression in CD8^+^ T cells from GBF for proteins involved in the granzyme-mediated programmed cell death signaling pathway and proteins derived from the MSigDB C5 ‘cell killing’ biological process gene set, ordered by log fold change between older (≥50 years) and younger (<50 years) donors. Pie charts in **d** show donor-level group means. Box plots show median with minimum-to-maximum range. *p* value in **b** were determined by two-tailed unpaired *t*-test. *p* value in **e** and **f** were determined by two-tailed unpaired *t*-test for KF and one-way ANOVA with Tukey’s multiple-comparisons test for GBF. PAGE, parametric analysis of gene set enrichment; T_RM_, tissue-resident memory T cell; NK, natural killer cell; ILC, innate lymphoid cell.

**Extended Data Figure 4.**
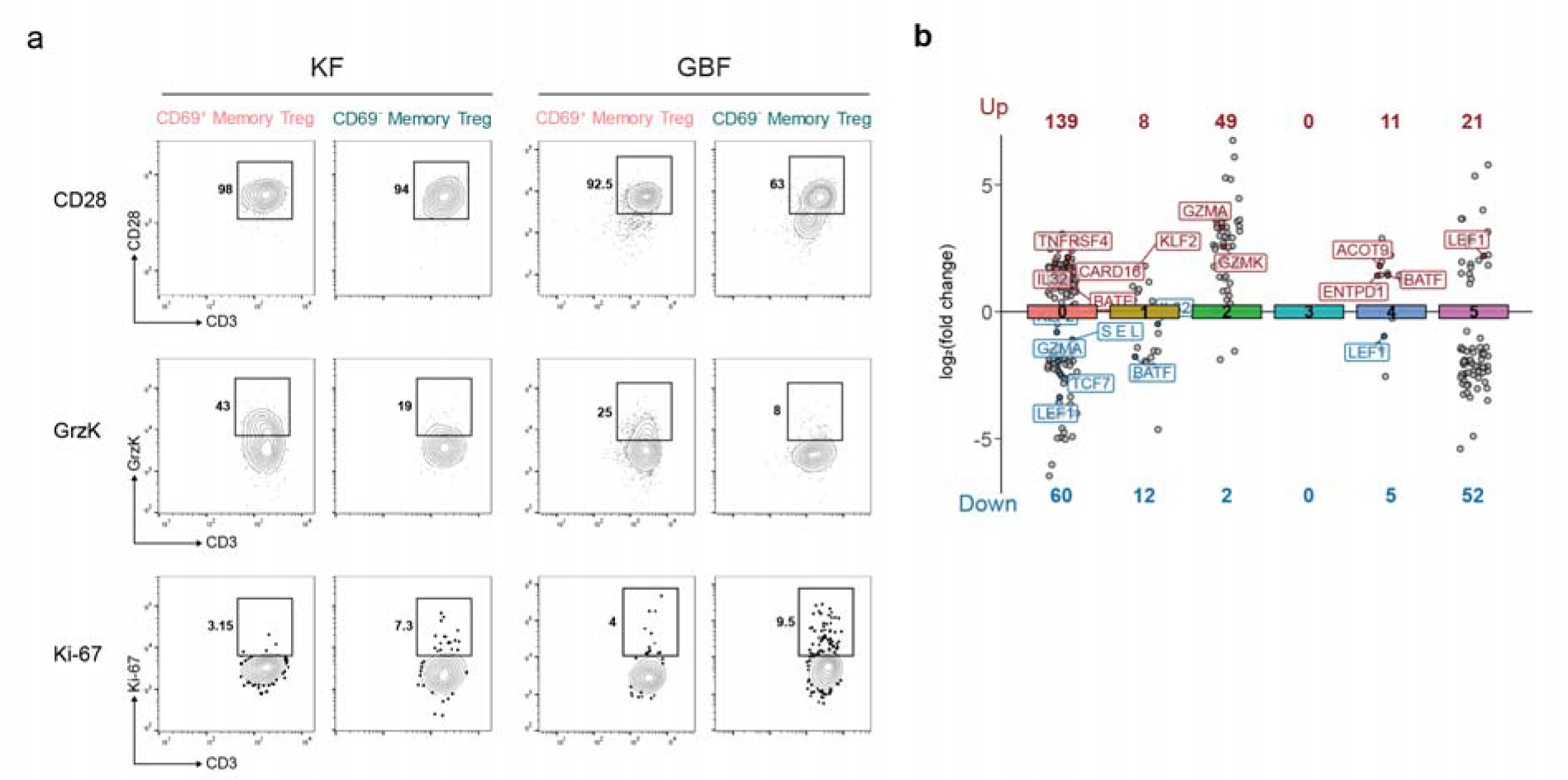
Treg phenotypic and transcriptional remodeling in aging VAT. a, Representative flow cytometry plots showing CD28, granzyme K and Ki-67 expression in CD69^+^ and CD69^-^ memory Tregs from KF and GBF. Memory Tregs were defined as CD4^+^Foxp3^+^CD45RO^+^ Tregs and further stratified by CD69 expression. Numbers indicate the frequencies of marker-positive cells within each gated subset. **b**, Marker genes distinguishing the six Treg subclusters (clusters 0–5, corresponding to Treg0–Treg5), each compared against all other cells. Points represent individual genes (adjusted *p* < 0.01), positioned by log fold change. The numbers above and below each cluster give the count of enriched (red) and depleted (blue) genes.

**Extended Data Figure 5.**
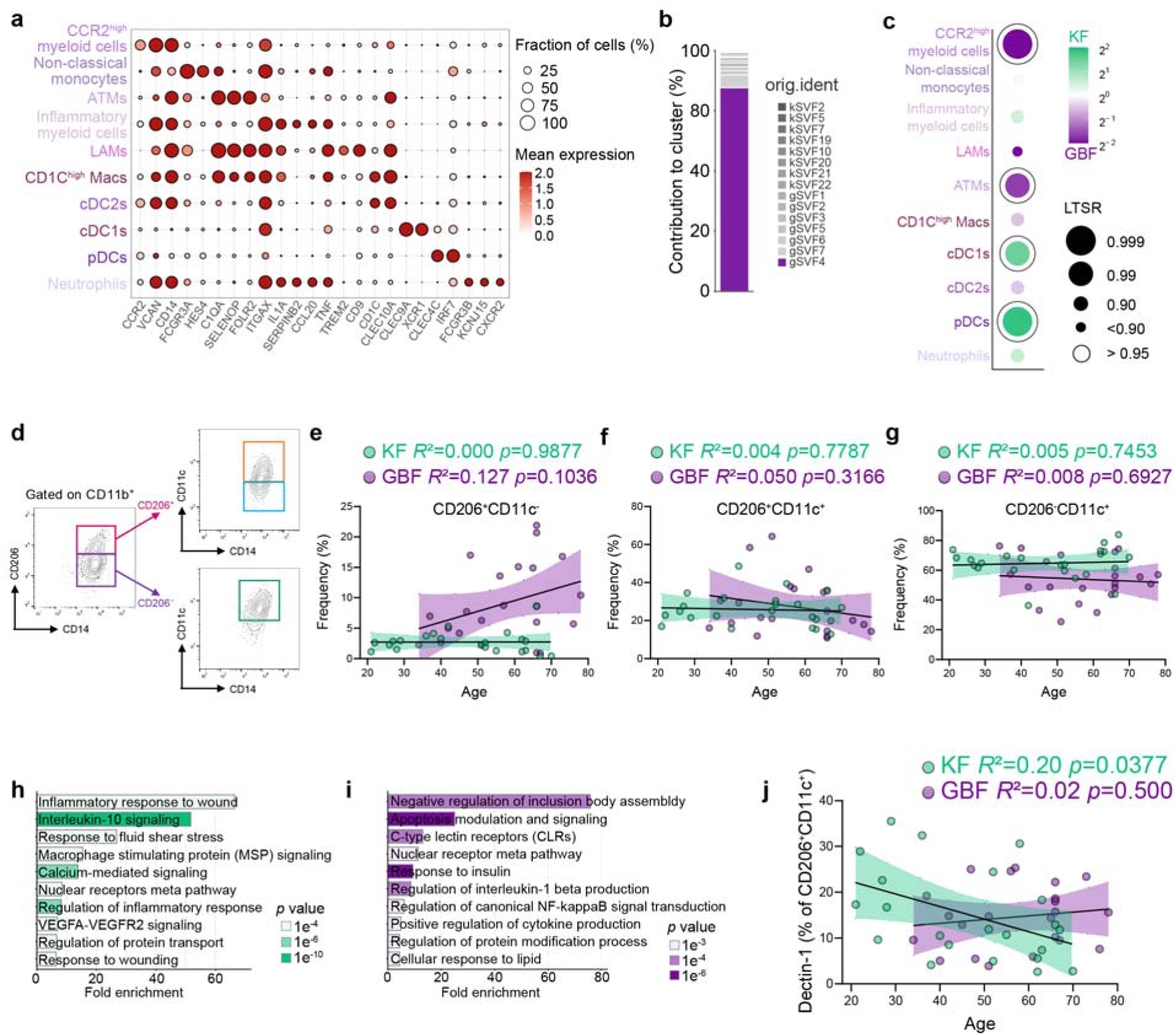
Additional validation of myeloid subsets and their depot-specific inflammatory programs. a, Dot plot of subcluster-defining marker genes across the myeloid subsets. Dot size encodes the fraction of expressing cells (%); color encodes mean expression. **b,** Stacked bar plot showing the donor-level contribution to *CCR2*^high^ myeloid cell cluster, colored by sample identity. **c,** Bubble plot of depot enrichment for each myeloid subset. Color encodes the KF-versus-GBF log2 fold change (green, KF; purple, GBF); bubble size encodes LTSR; open circles indicate LTSR > 0.95. **d,** Representative flow cytometry gating plots for CD11c^+^ myeloid cells in VAT. CD45^+^ leukocytes were gated, lymphocyte events were excluded, and CD11b^+^CD16^-^ myeloid cells were further divided into CD206^+^ and CD206^-^ subsets, followed by CD11c^+^ gating within each subset. **e–g,** Frequency of CD206^+^CD11c^-^ (**e**), CD206^+^CD11c^+^ (**f**), and CD206^-^CD11c^+^ (**g**) cells versus donor age in the KF (green) and GBF (purple). **h,i,** Metascape pathway enrichment analysis of differentially expressed genes upregulated in KF (**h**) and GBF (**i**) inflammatory myeloid cells. The top ten enriched terms are shown, ranked by statistical significance. Bar length indicates fold enrichment and bar shading indicates enrichment *p* value (color scale at right). **j,** Dectin-1^+^ cell frequency among CD206^+^CD11c^+^ cells versus donor age in the KF (green) and GBF (purple). For **e–g**, and **j**, *R*² and *p* value are indicated; lines, linear regression; shaded band, 95% confidence interval.

**Extended Data Figure 6.**
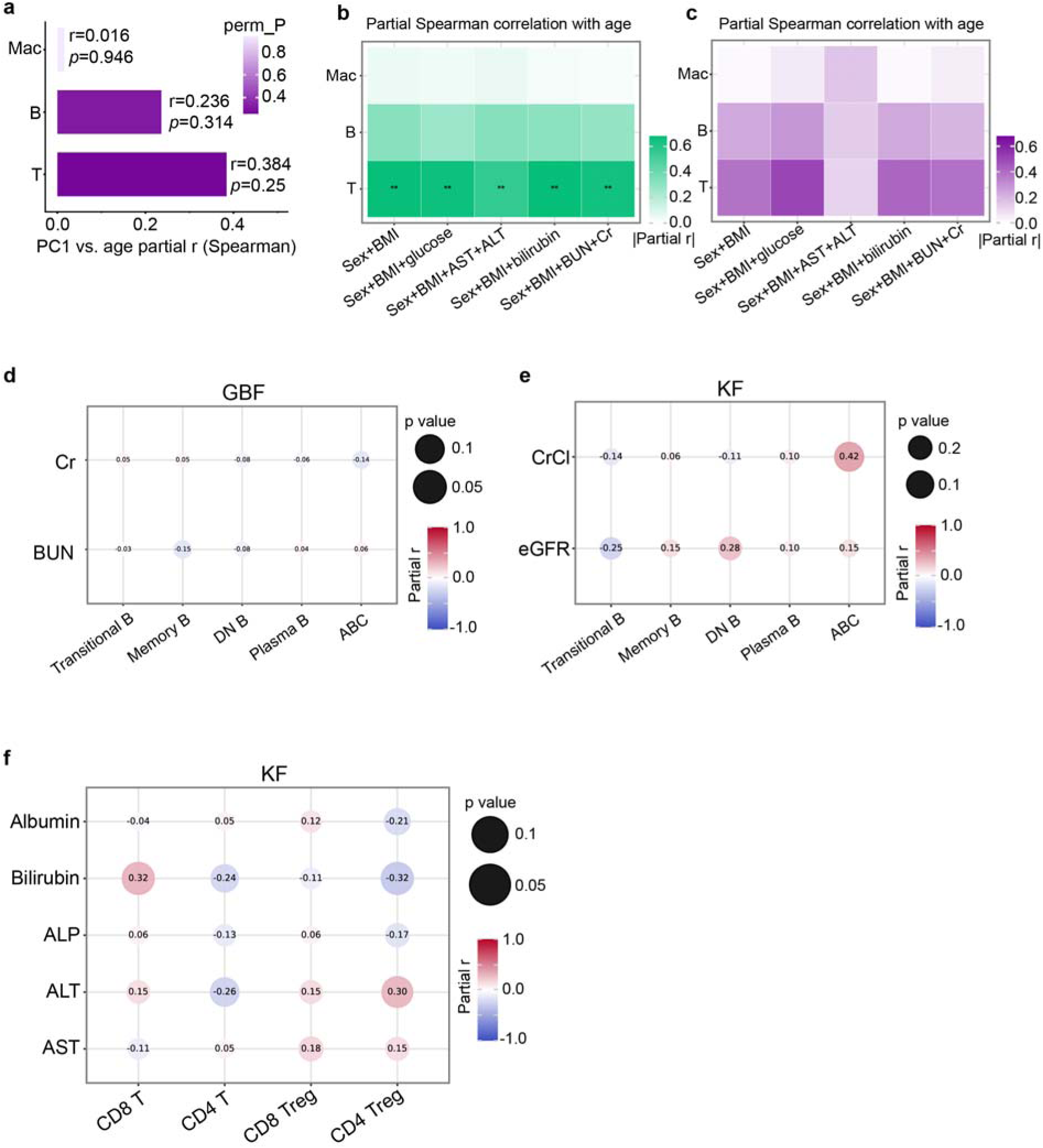
Robustness of age-associated T cell remodeling and depot-specific immune–organ associations. a, Partial Spearman correlation coefficients between each immune block PC1 and chronological age in the GBF, adjusted for sex and BMI. Color intensity encodes permutation-derived *p*-values (perm_P). **b,c,** Heatmaps of partial Spearman correlation coefficients between each immune block PC1 and chronological age in the KF **(b)** and GBF **(c)** depots, evaluated across five progressively expanded covariate adjustment sets. Color encodes absolute value of partial r. **d,** Bubble plot of feature-level pairwise partial Spearman correlations between B cell subsets and kidney biomarkers in the GBF depot, adjusting for sex and age. **e,** Bubble plot of partial Spearman correlations between B cell subsets and positive indicators of kidney function in the KF, adjusting for sex and age. **f,** Bubble plot of feature-level pairwise partial Spearman correlations between αβ T cell subsets and individual liver biomarkers in the KF depot, adjusting for sex and age. For all bubble plots, bubble size encodes *p*-value magnitude; color encodes signed partial r (red = positive, blue = negative). **q < 0.05.

