## Supplementary information for "The anatomical compartment defines distinct immune remodeling within human visceral adipose tissue during aging"

**Supplementary figures**

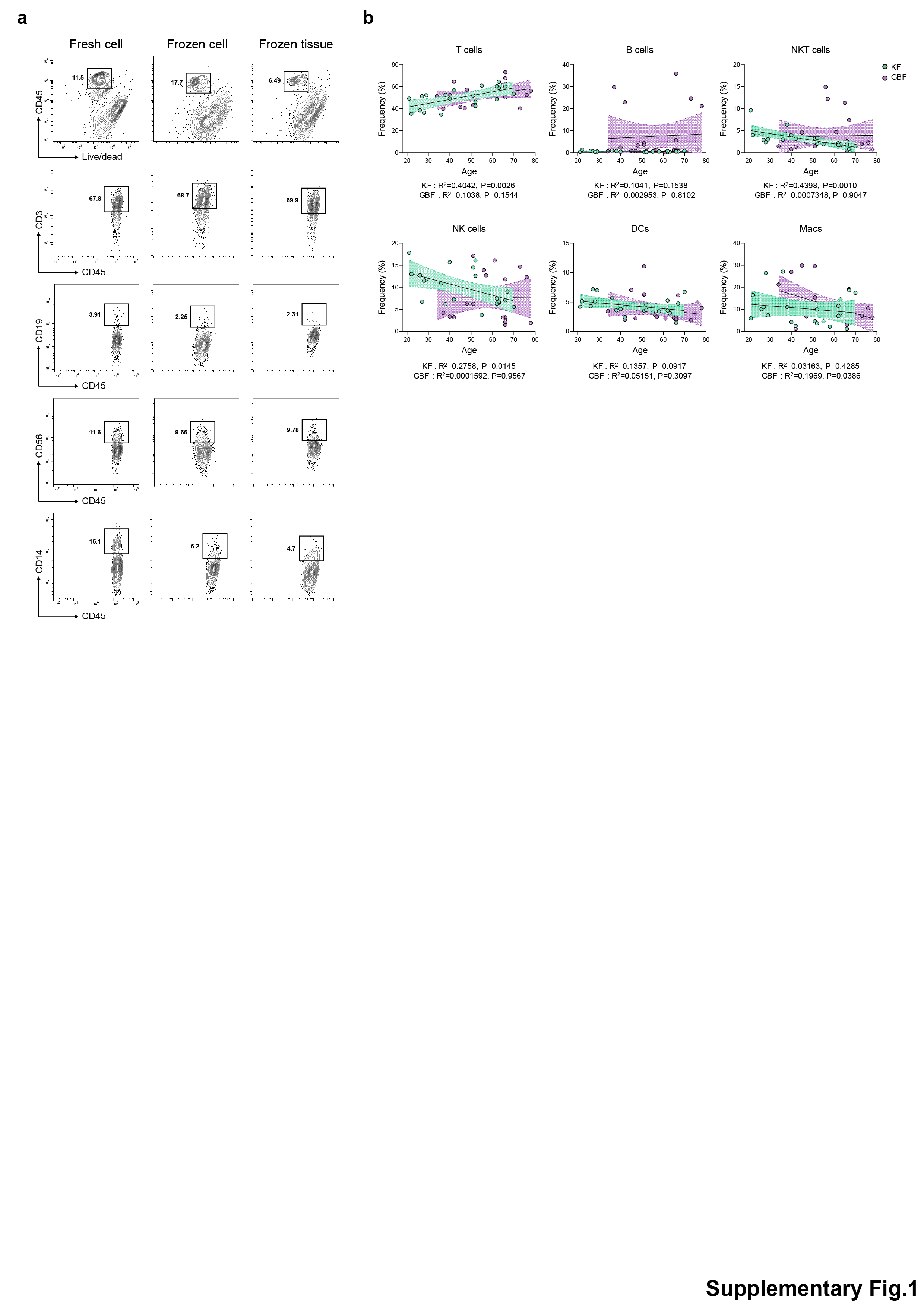

**Supplementary Figure 1. Validation of VAT sample preparation and age-associated immune cell changes.**

**a**, Representative flow cytometry plots comparing three sample preparation methods for VAT immune cell analysis: freshly processed cells, cryopreserved single-cell suspensions and cryopreserved tissue. **b**, Regression plots showing the frequencies (**b**) of the indicated CD45^+^ immune cell subsets as a function of donor age in the validation cohort (KF, n = 22; GBF, n = 22). Lines indicate simple linear regression fits; shaded areas indicate 95% confidence intervals. *R*² and *p* values were derived from simple linear regression.

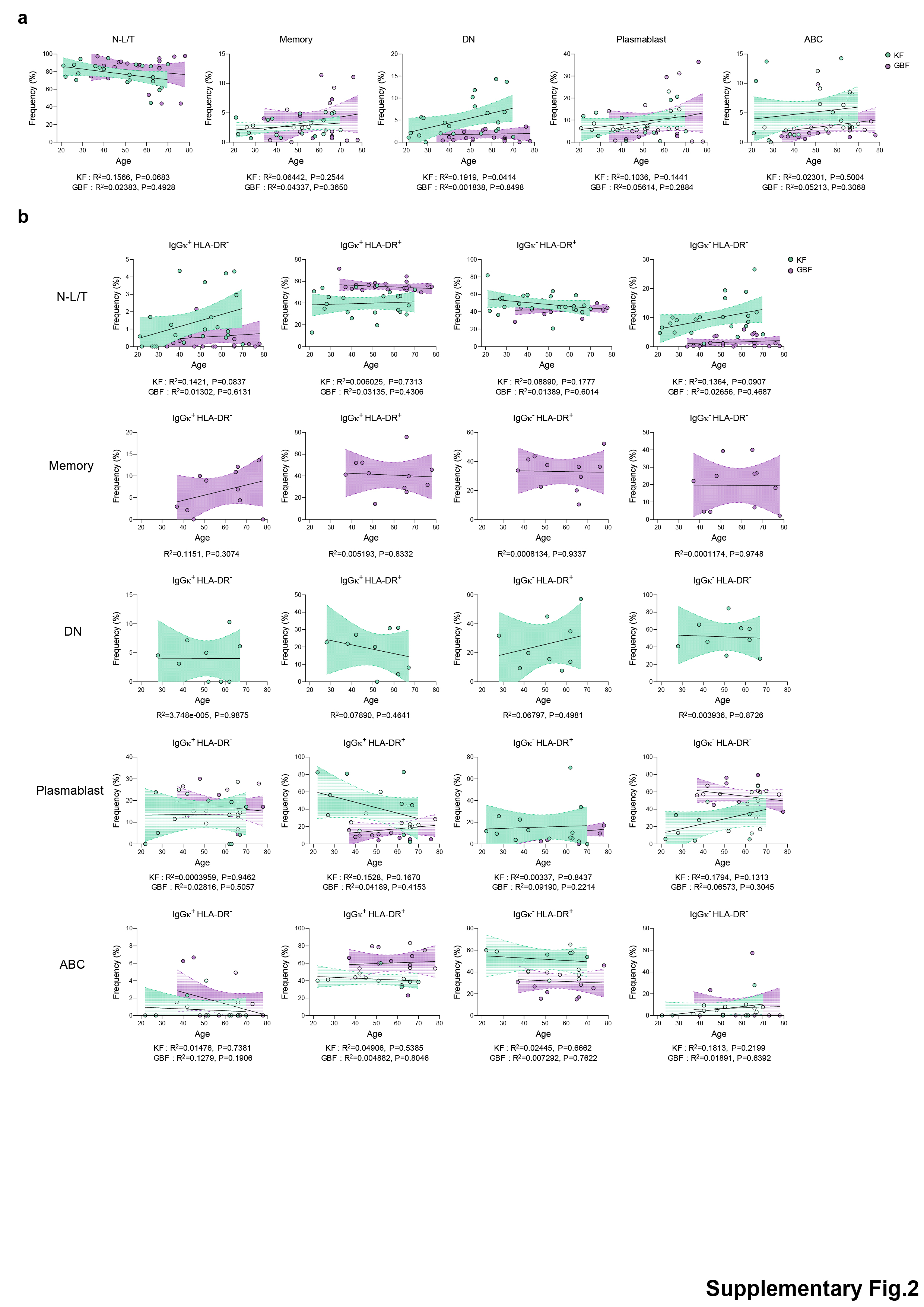

**Supplementary Figure 2. Age-associated B cell subset and activation-state changes in VAT.**

**a**,**b**, Regression plots showing the frequencies of the indicated CD19^+^ B cell subsets (**a**) and IgGκ/HLA-DR-defined states within each B cell subset (**b**) as a function of donor age in KF (n = 22) and GBF (n = 22). Lines indicate simple linear regression fits; shaded areas indicate 95% confidence intervals. *R*² and *p* values were derived from simple linear regression.

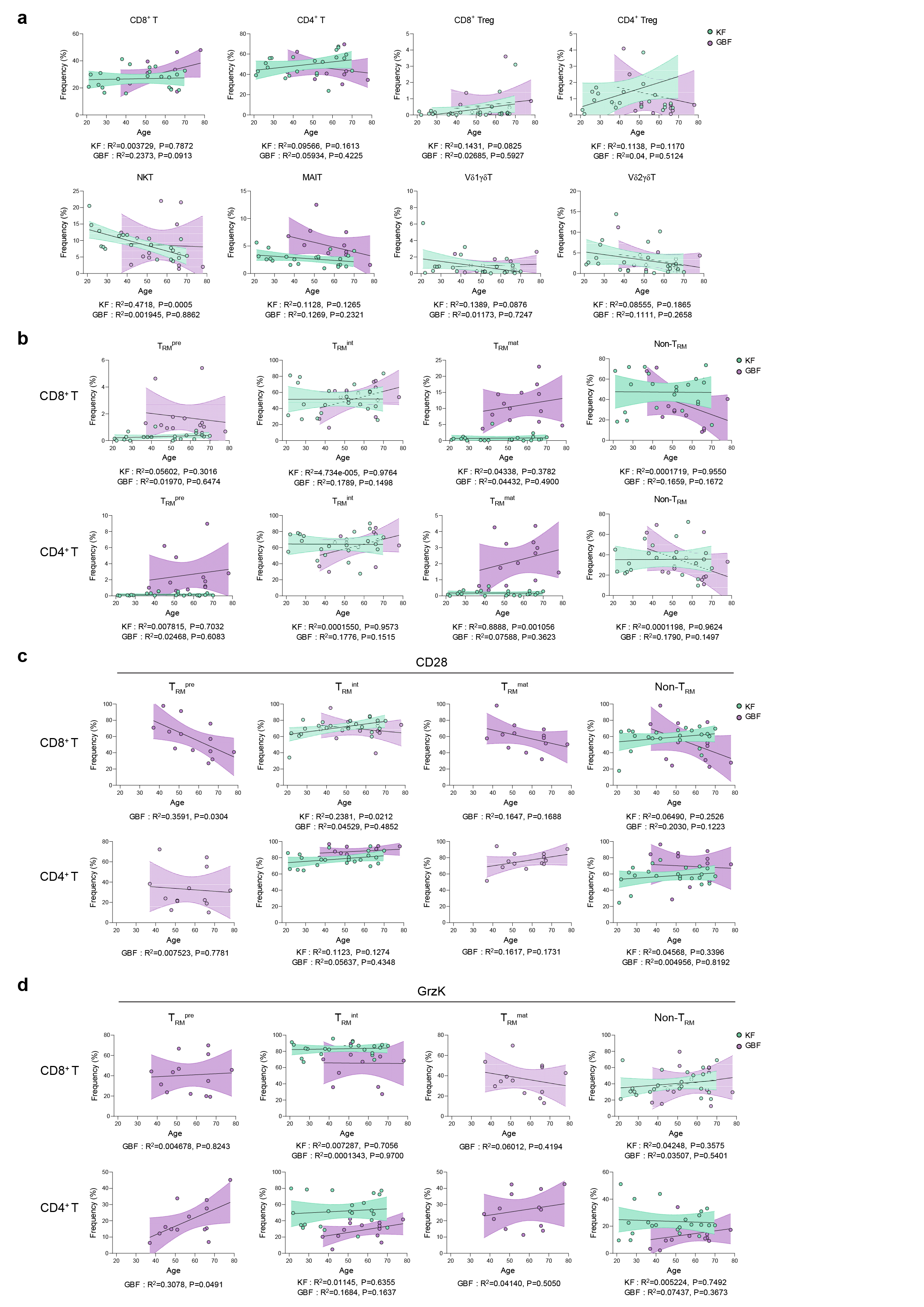

**Supplementary Figure 3. Age-associated T cell and T_RM_-related phenotypic changes in VAT.**

**a-d,** Regression plots showing the frequencies of the indicated CD3^+^ T cell subsets (**a**), CD69/CD103-defined T_RM_-related states (**b**), CD28^+^ cells within each T_RM_-related state (**c**) and granzyme K^+^ cells within each T_RM_-related state (**d**) as a function of donor age in KF (n = 22) and GBF (n = 13). Lines indicate simple linear regression fits; shaded areas indicate 95% confidence intervals. *R*² and *p* values were derived from simple linear regression. T_RM_, tissue-resident memory.

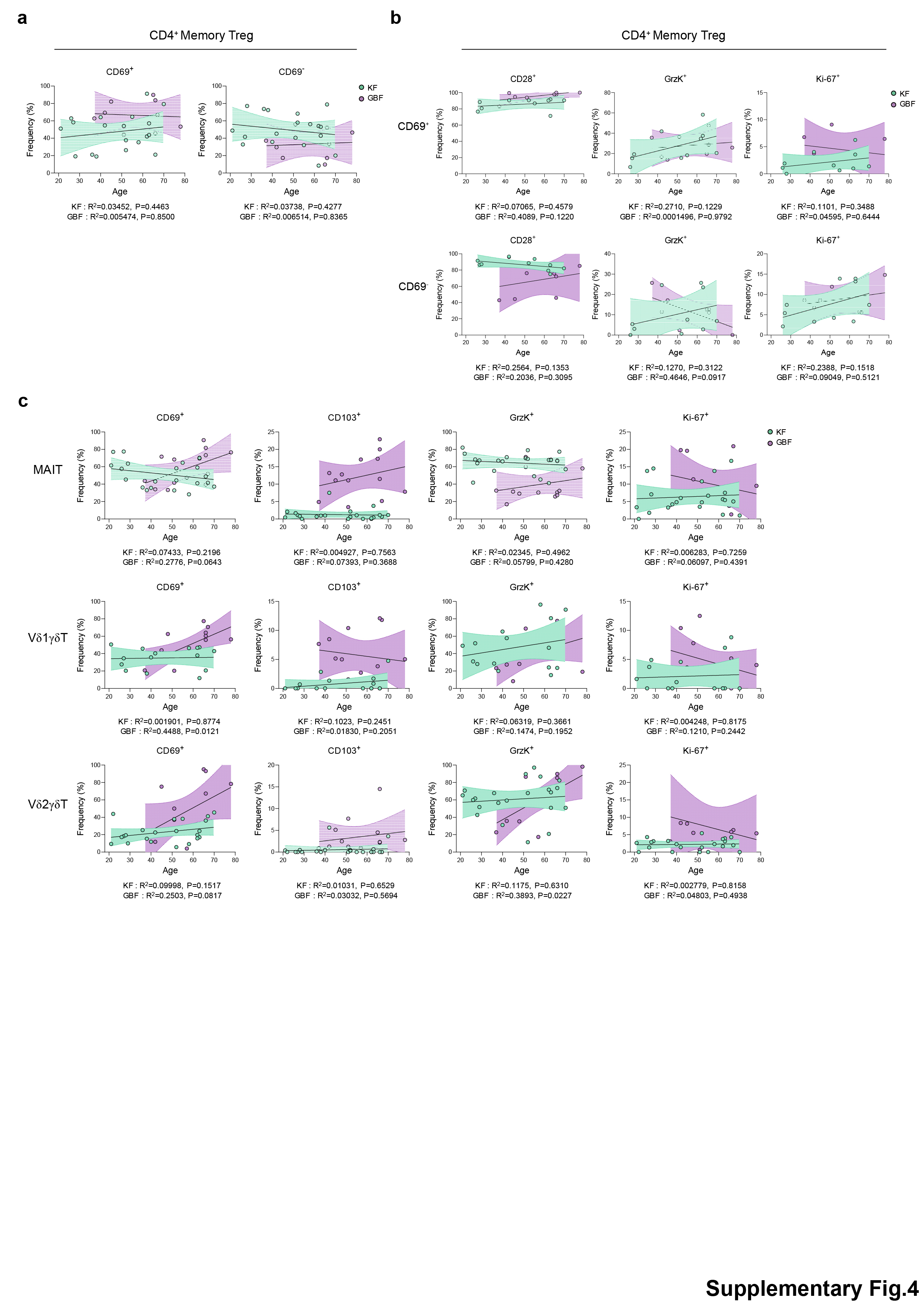

**Supplementary Figure 4. Age-associated phenotypic remodeling of memory Tregs and innate-like T cells in VAT.**

**a-c**, Regression plots showing the frequencies of CD69^+^ and CD69^-^ subsets among CD4^+^ memory Tregs (**a**), CD28^+^, granzyme K^+^ and Ki-67^+^ cells among CD69^+^ and CD69^-^ CD4^+^ memory Treg subsets (**b**), and CD69^+^, CD103^+^, granzyme K^+^ and Ki-67^+^ cells among MAIT cells and Vδ1^+^ and Vδ2^+^ γδ T cells (**c**) as a function of donor age in KF and GBF. Lines indicate simple linear regression fits; shaded areas indicate 95% confidence intervals. *R*² and *p* values were derived from simple linear regression. Treg, regulatory T cell; MAIT, mucosal-associated invariant T cell.
